# Computational insights on the destabilizing mutations in the binding site of 3CL-protease SARS-CoV-2 Omicron (VOC)

**DOI:** 10.1101/2023.05.24.542061

**Authors:** Samee Ullah, Afreenish Amir, Aamer Ikram, Caterina Vicidomini, Rosanna Palumbo, Giovanni N. Roviello

## Abstract

The COVID-19 (Corona Virus Disease 19) pandemic, caused by Severe Acute Respiratory Syndrome Coronavirus 2 (SARS-CoV-2), is causing enormous difficulties in the world’s economies and there is uncertainty as to whether the current prophylactic measures will offer adequate protection globally after the appearance of virus variants that like that indicated as Omicron emerged in the presence of global vaccine-based immunization. While several studies are available describing the main differences in the spike protein of Omicron compared to the other variants previously emerged, there was no structural insights into the 3CL-protease (3CL^pro^) associated to the new variant. Herein, we performed a computational study based on genomic data and amino acid sequences available in the most updated COVID-19-related databases that allowed us to build up *in silico* the 3D structure of Omicron 3CL^pro^. Moreover, by molecular dynamics simulation we demonstrated that currently available drugs acting as inhibitors of the SARS-CoV-2 main protease could be less effective in the case of Omicron variant due to the different chemical interactions in the binding site occurred after the recent amino acid mutations. Ultimately, our study highlights the need of exploiting *in silico* and in vitro methods to discover novel 3CL^pro^ inhibitors starting from the computationally based structure we presented herein, and more in general to direct the major efforts to targeting the most conserved 3CL^pro^ regions that appeared unchanged in the context of the Omicron variant.

## Introduction

During 2020 the world witnessed the spread of severe acute respiratory syndrome coronavirus 2 (SARS-CoV-2), which led to the potentially deathly coronavirus disease 2019 (COVID-19)[1]. SARS-CoV-2 appears close to both the severe acute respiratory syndrome (SARS) and the Middle East respiratory syndrome (MERS) through its similar immunopathology[1]. Originating in Wuhan, China, COVID-19 rapidly succeeded in spreading around the globe, resulting in the current pandemic. To date, there have been more than 6,088,152 deaths associated to the disease globally (https://www.worldometers.info/coronavirus/ accessed on 18 March 2022). COVID-19 has also forced numerous countries to face difficult sanitary conditions with overcrowded hospitals, becoming, thus, the cause of great instability in the world from both a sanitary and socio-economic perspective [2–5]. While many are hopeful to have found in the ongoing global mass immunization using the already developed anti-COVID-19 vaccines [6, 7] a mid-term valid solution [6], new SARS-CoV-2 variants are periodically emerging and there is uncertainty as to whether or not the current immunization will offer adequate protection [8]. Until mass vaccination has proven successful, many research groups continue to devote their efforts to the exploration of more immediate solutions to the infection. One such strategy is the trial of pharmaceuticals that are already known to be effective in the treatment of other viral and COVID-19-related disorders [9] and is often indicated as ‘drug repurposing’ [9–12], but other strategies are also being explored such as herbal medicine [13] and other recently-proposed approaches [14, 15] in the therapy of SARS-CoV-2 [13, 16]. Of course looking into ways of preventing further pandemic outbreaks and strengthening human defenses against zoonoses and viral diseases is also of great importance[17, 18] but anti-COVID-19 therapy [10, 19] is the only way to respond to the urgency of saving lives in the hospitals. In this context, many anti-COVID-19 drugs being tested in silico and experimentally are inhibitors of the 3CL-protease of SARS-CoV-2 (3CL^pro^), that represents one of the most attractive biomolecular targets for COVID-19 therapy due the lack of its homologs in humans, as well its crucial role in viral replication [20].

Although the human coronaviruses have genomes less variable than other viruses, the receptor binding domain (RBD) contained in the spike protein is the most mutable region, with mutation sites that may lead to worrying antibody escape variants [21–23]. Several mutations of SARS-CoV-2 involving the spike protein have led to reduced antibody binding and increased infectivity [24]. Hence, the current concern on the new antibody-escaping mutations emerging in the context of the current mass immunization based on spike-targeting vaccines. In fact, these variants could make the current anti-COVID-19 protection less effective and the process of new vaccine development even more complicated.

In late November 2021, after the alpha, beta, and delta SARS-CoV-2 variants of concern (VOC), a new one, the VOC Omicron, emerged in a context in which vaccine and not only natural immunization was common[25]. While VOC Omicron was first identified by scientists in South Africa, only few days after its announcement by WHO, a number of cases of Omicron have been recorded in numerous countries such as Austria, Belgium, Canada, United Kingdom, Denmark, France, Germany, and Italy[26] testifying the extremely rapid spread of the virus over the globe.

While several studies are available in the scientific literature describing the main differences in the spike protein of VOC Omicron compared to the previously-detected SARS-CoV-2 variants[25, 27], there is no structural insights into the 3CL-protease (3CL^pro^) associated to the new variant. Aiming at opening the way to new anti-COVID-19 drug development, in this work we performed a computational study starting from genomic data and crystal structure available in the most updated COVID-19-related databases for the VOC, that allowed us to build up *in silico* the 3D structure of the Omicron 3CL^pro^. Moreover, by molecular dynamics simulation we investigated the potential inhibitory potential of the currently available drug Nirmatrelvir, that acts as an inhibitor of the SARS-CoV-2 main protease[28], to estimate its effectiveness in the case of Omicron variant after the several amino acid mutations revealed.

## Materials and Methods

The complete workflow of the present study is displayed in figure 1.

**Figure 1.**
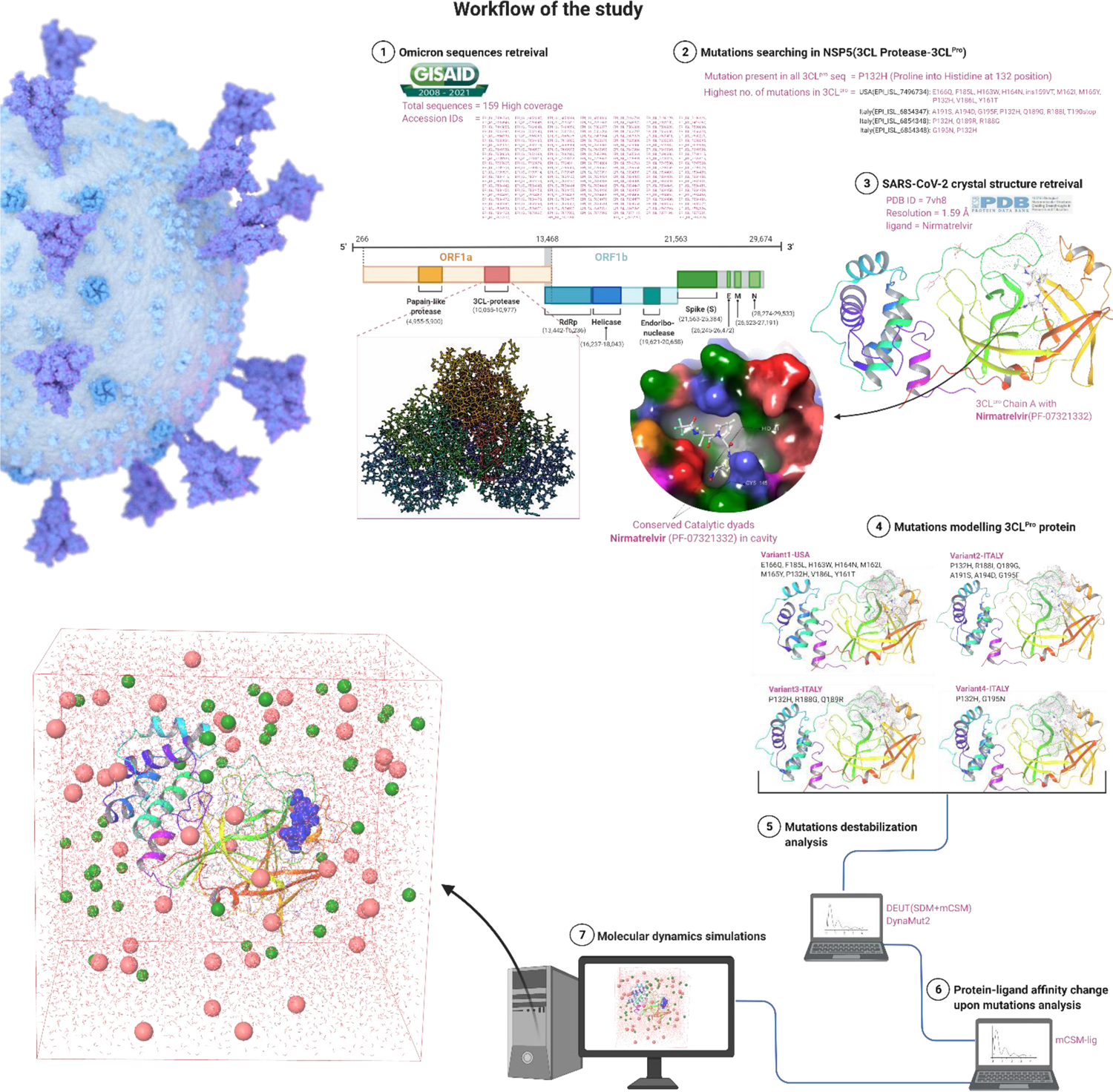
Workflow of the current study

### Sequence retrieval and structure modelling of M^pro^ Omicron VOC

The GISAID database EpiCoV^TM^ searched for the NSP5 sequences of SARS-CoV-2 Omicron variant of concern (VOC) [29]. The mutations reported in the non-structural protein (NSP5) which encodes main protease enzyme (M^pro^) vital for viral replication were mined with the following Accession IDs in supply table1. The mutations were modelled in a crystal structure PDB ID: 7VH8 complex with its inhibitor Nirmatrelvir (oral antiviral developed by Pfizer).

In this study, the crystal structure of the SARS-CoV-2 main-protease enzyme (PDB ID: 7VH8) was obtained with a resolution of 1.59Å from the Research Collaboratory for Structural Bioinformatics Protein Data Bank (RCSB-PDB) (https://www.rcsb.org/) [30, 31]. Four separate 3D structures of the main protease were used to incorporate mutations, including variant1 (P132H, Y161T, M162I, H163W, H164N, M165Y, E166Q, F185L, V186L) reported from the USA and variant2, variant3-4 (P132H, RI88I, Q189G, A191S, A194D, G195F), (P132H, R188G, Q189R), and (P132H, G195N) reported from Italy, respectively. The M^pro^ three-dimensional crystal structure was imported into the maestro visualizer and the mutations were incorporated at each position via the mutation build panel. Hydrogen atoms were added and optimized using the optimization panel and a default minimization of 0.2 Å RMSD was achieved before simulations using the OPSL3e force field [32–34].

### Molecular dynamic simulation protocol

The M^pro^ structures variants 1-4 were evaluated using the Desmond simulation engine for the 100ns simulation period [35]. The simulations were conducted using an AMD Ryzen^TM^ 7 5800H processor with 64-GB of RAM and Linux Fedora 37 Scientific (64bit) operating system (OS).

### System building

A solvent model (simple point charge-SPC) using the default parameters of the OPLS3e forcefield was constructed for all 4 Variants of the 3CL protease. The boundary conditions for the simulation box were set to cubic at a 10 Å distance in x, y, and z coordinates, with a calculated volume of 539852 Å3 for each case. A 0.1M salt concentration was added to the system, and the full system was neutralized by adding 5 Na ions after recalculating the necessary ions. The final system consisted of 52279 atoms.

### Trajectory production

For trajectory production, the full simulation systems of Variants 1-4 of the 3CL protease were imported into the Desmond molecular dynamics wizard. A simulation time of 100 ns was set for each system, with trajectories recorded at 100 ps intervals and approximately 1000 frames. The default temperature of 300 K and pressure of 1 atm were applied, with the NPT ensemble class used for all variants. The full systems were relaxed before simulation using the default “relax model before simulation” algorithm and the OPLS3e forcefield.

### M^pro^ mutations stability analysis

The structures of 3CL protease Variants1-4 were imported into DEUT (https://doi.org/10.1093/nar/gku411) algorithm webserver which uses two additional separate techniques namely SDM (https://dx.doi.org/10.1093%2Fnar%2Fgkr363) and mCSM (https://doi.org/10.1093/protein/10.1.7) and combines the prediction results via Support Vector Machines (SVM) to analyze the impact of the mutations on protein stability. The SDM approach calculates the difference in free energy between wild type (WT) and mutant (Mu) proteins by comparing the amino acid propensities in both the folded and unfolded states. Furthermore, in additional, to further support the mutational stability analysis, the structures were also evaluated through the DynaMut2 algorithm (https://doi.org/10.1002/pro.3942) which predicts the stability and flexibility of single and multiple point missense mutations.

### Protein-ligand affinity change upon mutations analysis

The Variants1-4 of having 3CL protease-Nirmatrelvir complexes were analyzed for the protein-ligand affinity change upon mutations. For the affinity, the Ki 3.11nM of Nirmatrelvir used and the protein-ligand affinity change upon mutations predicted.

## Results

### M^pro^ sequence retrieval and structure modelling of mutations

This study focused on 58 high coverage sequences that reported mutations in the NSP5 (3CL-protease) gene. One significant mutation was identified, the P132H (Proline to Histidine substitution at position 132), which is a conserved amino acid change in the 3CL-pro Omicron VOC. The comparison of 3CL-protease mutations between Omicron and previously reported SARS-CoV-2 variants showed critical differences, as illustrated in table 1 (cite source).

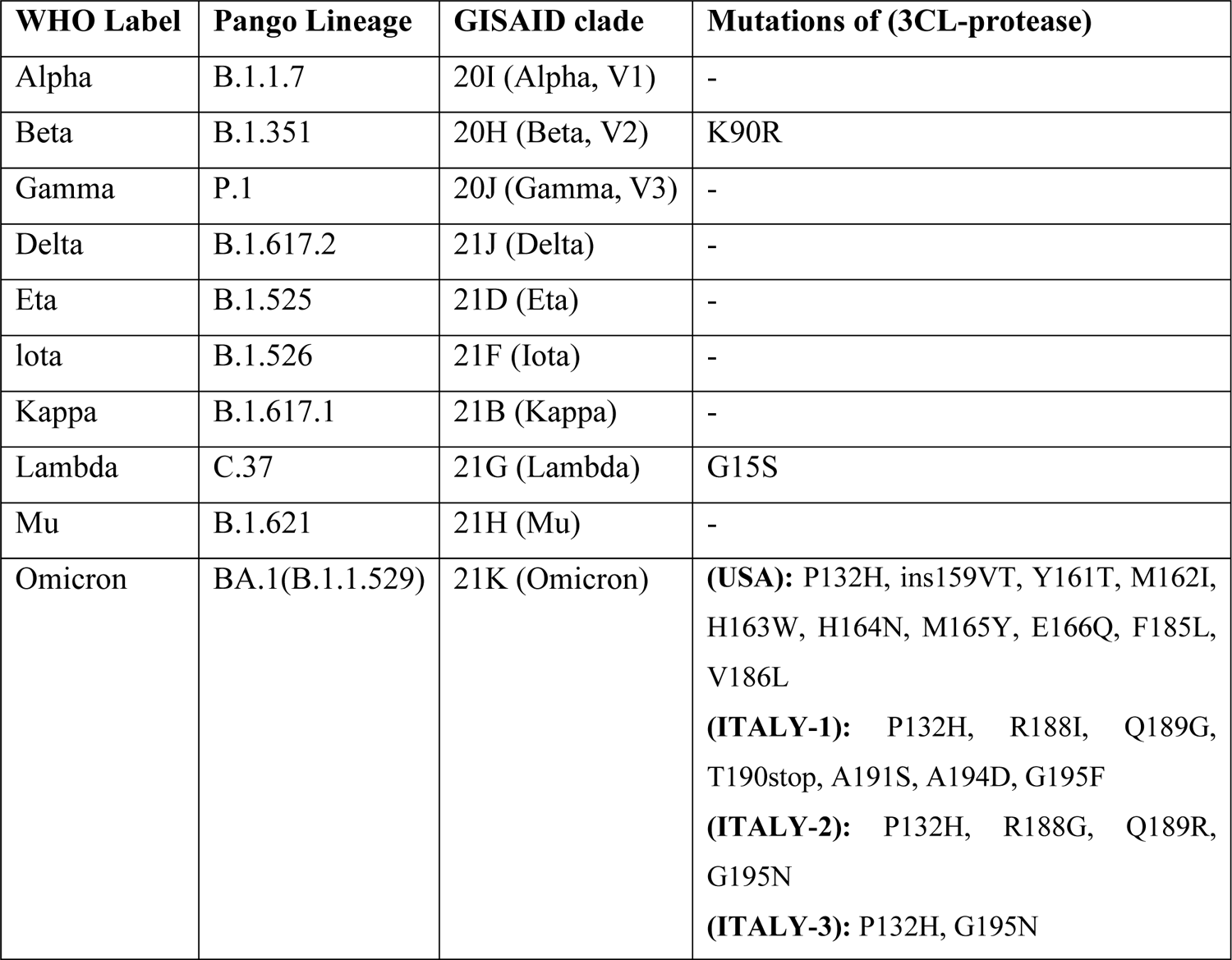

A 52-Year-old male patient in the USA has reported the highest number of amino acid mutations (P132H, ins159VT, Y161T, M162I, H163W, H164N, M165Y, E166Q, F185L, V186L) in the 3CL-protease Omicron sequences. Meanwhile, the second to fourth highest number of mutations have been found in 8-Year-old female, 81-Year-old female, and 77-Year-old female from Italy respectively with mutations such as (P132H, R188I, Q189G, T190stop, A191S, A194D, G195F), (P132H, R188G, Q189R), and (P132H, G195N) as shown in table-S1.

To better understand the impact of critical mutations in the 3CL-protease of Omicron VOC, we utilized Insilco techniques to model the mutations into four 3CL-protease variants in the crystal structure PDB ID: 7VH8 (3CL-protease-Nirmatrelvir complex). Our goal was to determine if these mutations would result in the destabilization of the binding of small molecules in the active site or folding of the protease. Based on data from GISAID and RCSB, we excluded substitutions such as ins159VT and T190stop from our study. Our results, shown in figure 3A-D, demonstrated the impact of these critical mutations on the four 3CL-protease variants.

**Figure 3A-D:**
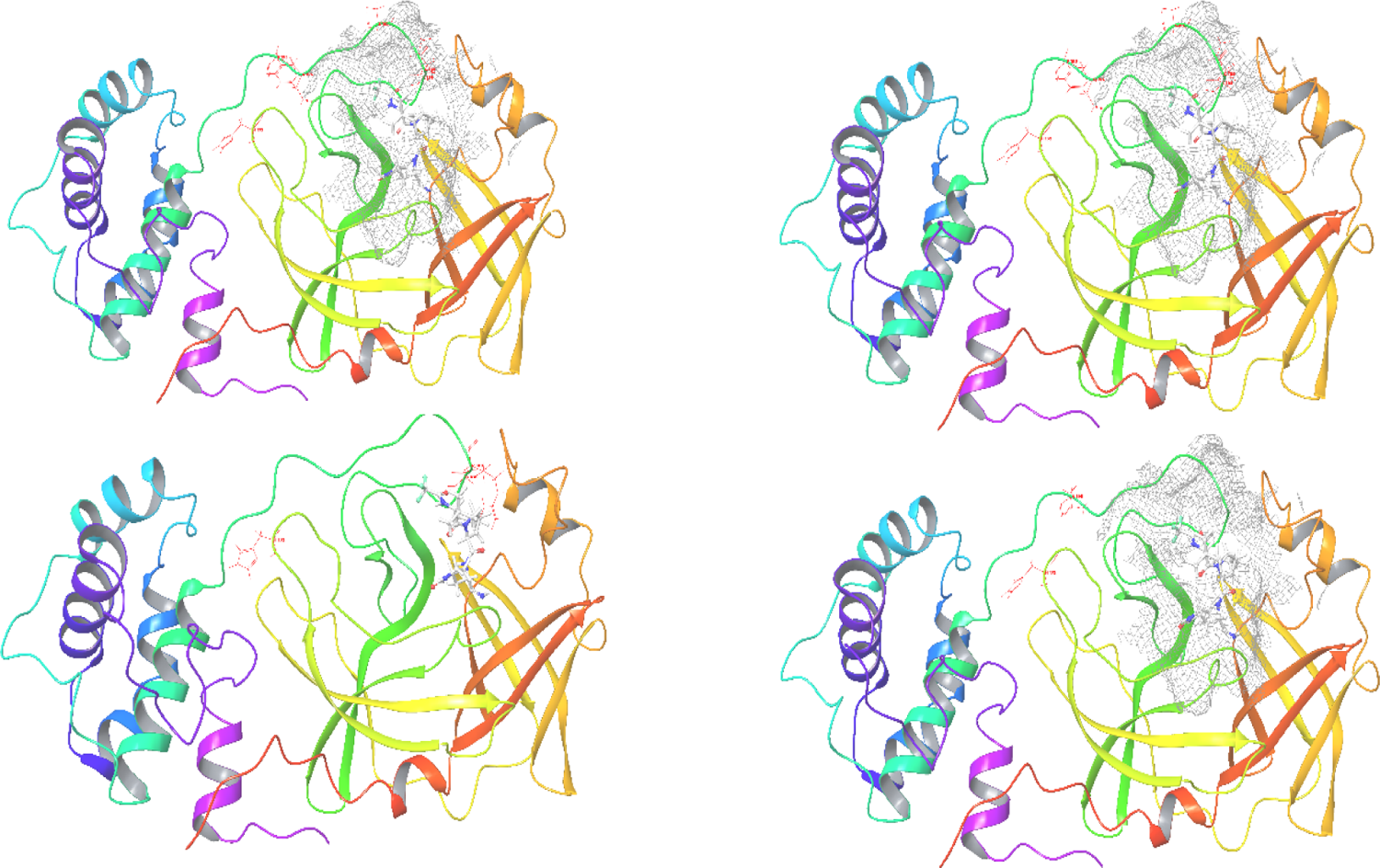
The red sticks symbolize mutations, and the grey mesh signifies the active site region of 3CL-protease Omicron VOC. Nirmatrelvir, shown in white element representation, serves as the native ligand.

### Critical mutations in the M^pro^ active site Omicron VOC

Previously two mutations K90R in Beta (B.1.351) VOC and G15S in lambda (C.37) reported in the 3CL-protease of SARS-CoV-2 variants (source). The analysis of Variant1, which has the highest number of mutations in the 3CL-protease, revealed that four mutations (H163W, H164N, M165Y, and E166Q) were located in the active site, while two mutations (Y161T and M162I) and two others (F185L and V186L) were located close to the pocket. In the analysis of Variant2, three mutations (R188I, Q189G, and A191S) were found in the pocket and two mutations (A194D and G195F) were located close to the pocket. Similarly, in Variant3-4, two mutations (R188G and Q189R) were found in the pocket and one mutation (G195N) was located close to the pocket. Interestingly, we found that the mutation P132H was present in all sequences analyzed, suggesting that it is a highly conserved change in the Omicron VOC 3CL-protease. This mutation was located at a distance of 17 Å from Nirmatrelvir in the active site.

We analyzed the recently reported critical mutations in the active site of the 3CL-protease Omicron VOC at protein three-dimensional structure level. The Variant1-USA (highest no. of mutations in 3CL-protease) revealed the (H163W: histidine into tryptophan at 163 positions, H164N: histidine into asparagine at 164, M165Y: methionine into tyrosine at 165 and E166Q: glutamate into glutamine at 166) are in the active site and the Y161T: tyrosine into threonine at position 161, M162I: methionine into isoleucine at 1621, F185L: phenylalanine into leucine at 185 and V186L: valine into leucine at 186 close to the pocket in figure 4-A. The Variant2-Italy 3CL-protease structures analysis revealed the (R188I: arginine into isoleucine at 188 positions, Q189G: glutamine into glycine at 189, A191S: alanine into serine at 191) in the pocket and the (A194D: alanine into aspartic acid at 194, G195F: glycine into phenylalanine at 195) close to the pocket in figure 4B. The Variant3-4Italy revealed the (R188G: arginine into glycine at 188 positions, Q189R: glutamine into arginine at 189) residues in the pocket and G195N: glycine into asparagine at 195 closes to the pocket in figure 4C-D. Interestingly, the mutation (P132H: proline into histidine at position 132) is found in all sequences suggesting a very conserved change adopted by Omicron VOC 3CL-protease far 17 Å distance from the Nirmatrelvir in the active site.

**Figure 4A.**
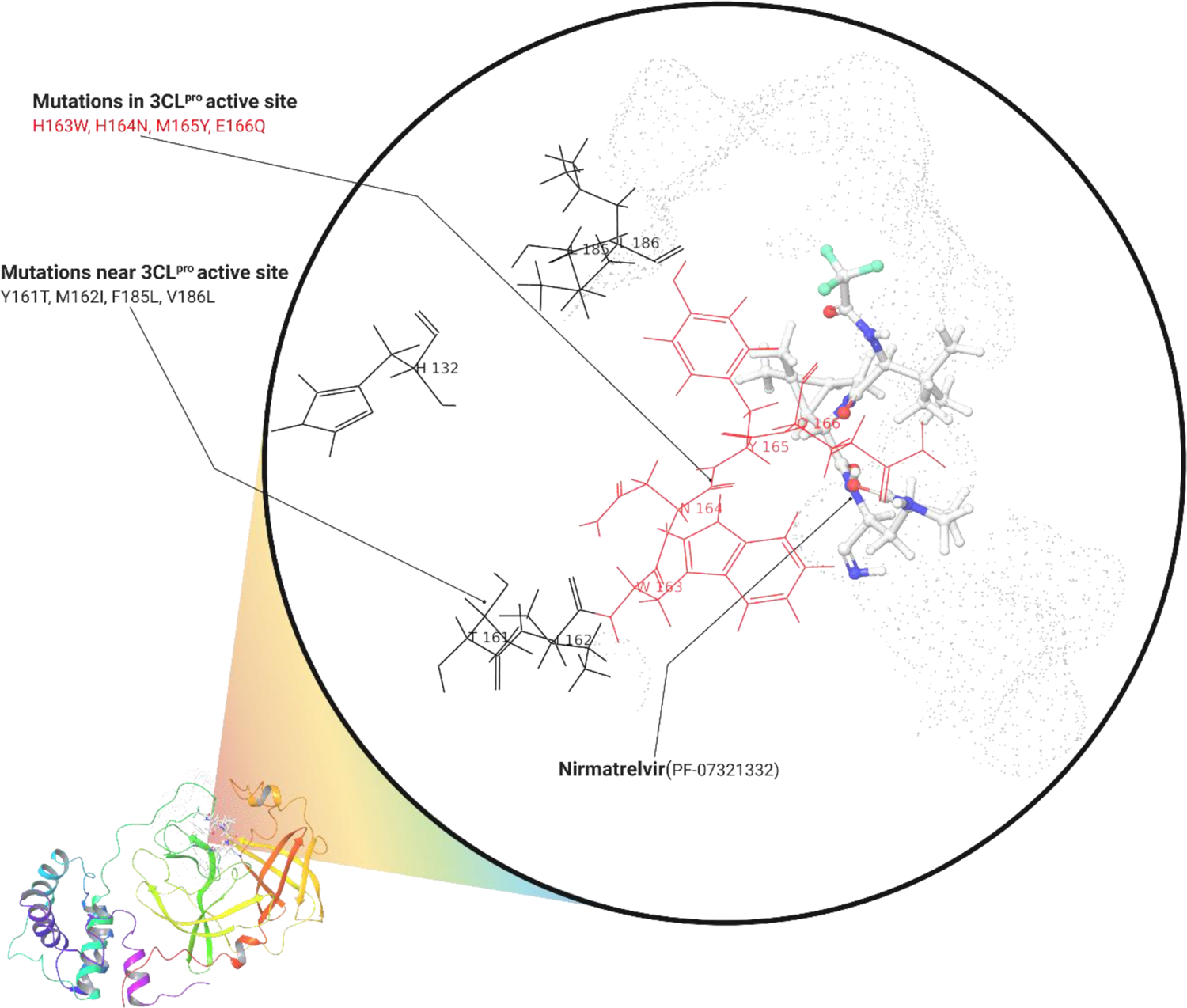
The 3CL-protease Variant1-USA (highest no. of mutations), with grey dots represent the active site region with bind Nirmatrelvir. The red colored sticks represent the mutations in the pocket and the black sticks close to the pocket except the P132H far.

**Figure 4B.**
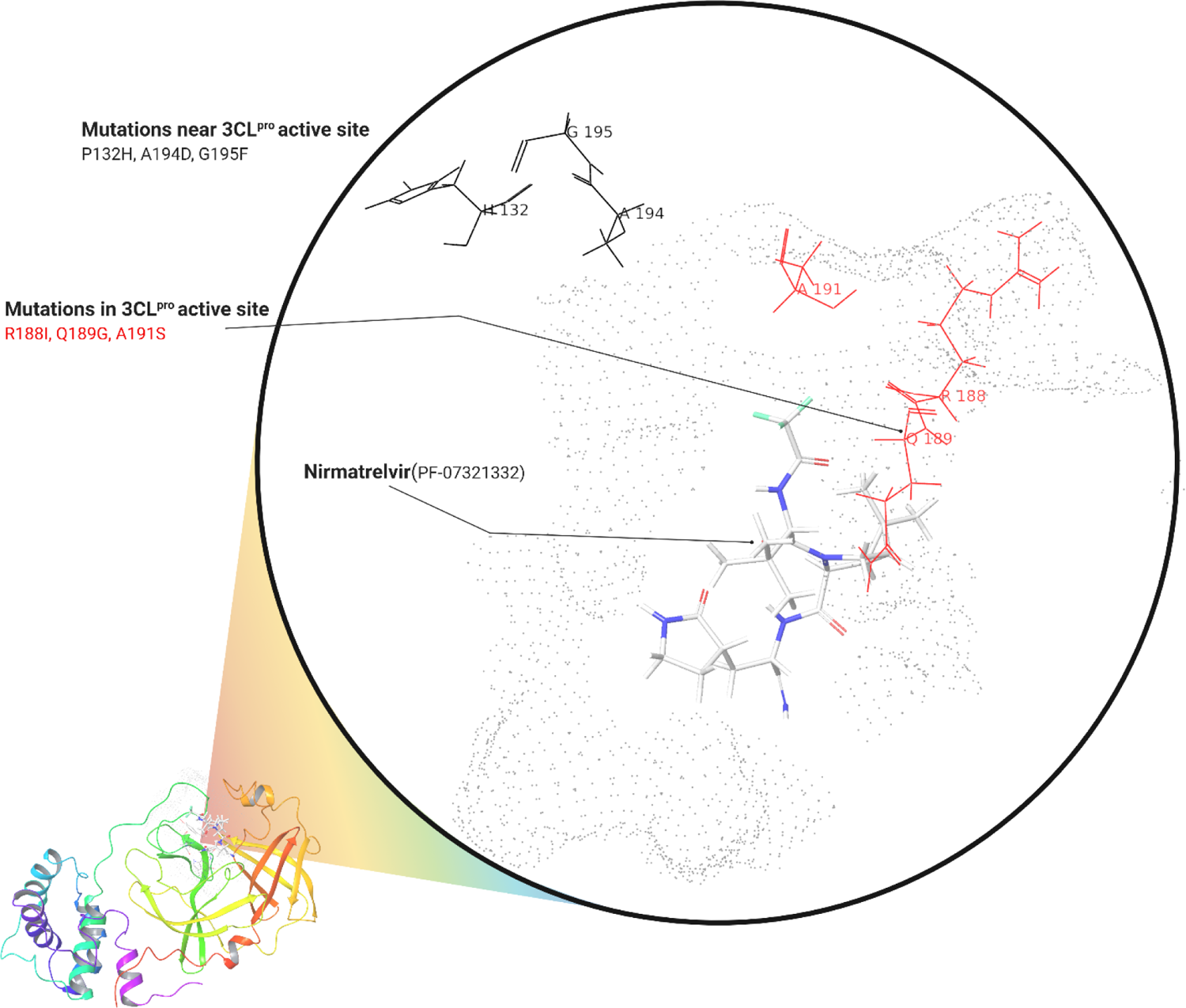
The 3CL-protease Variant2-Italy (2^nd^ highest no. of mutations), with grey dots represent the active site region with bind Nirmatrelvir. The red colored sticks represent the mutations in the pocket and the black sticks close to the pocket except the P132H far.

**Figure 4C.**
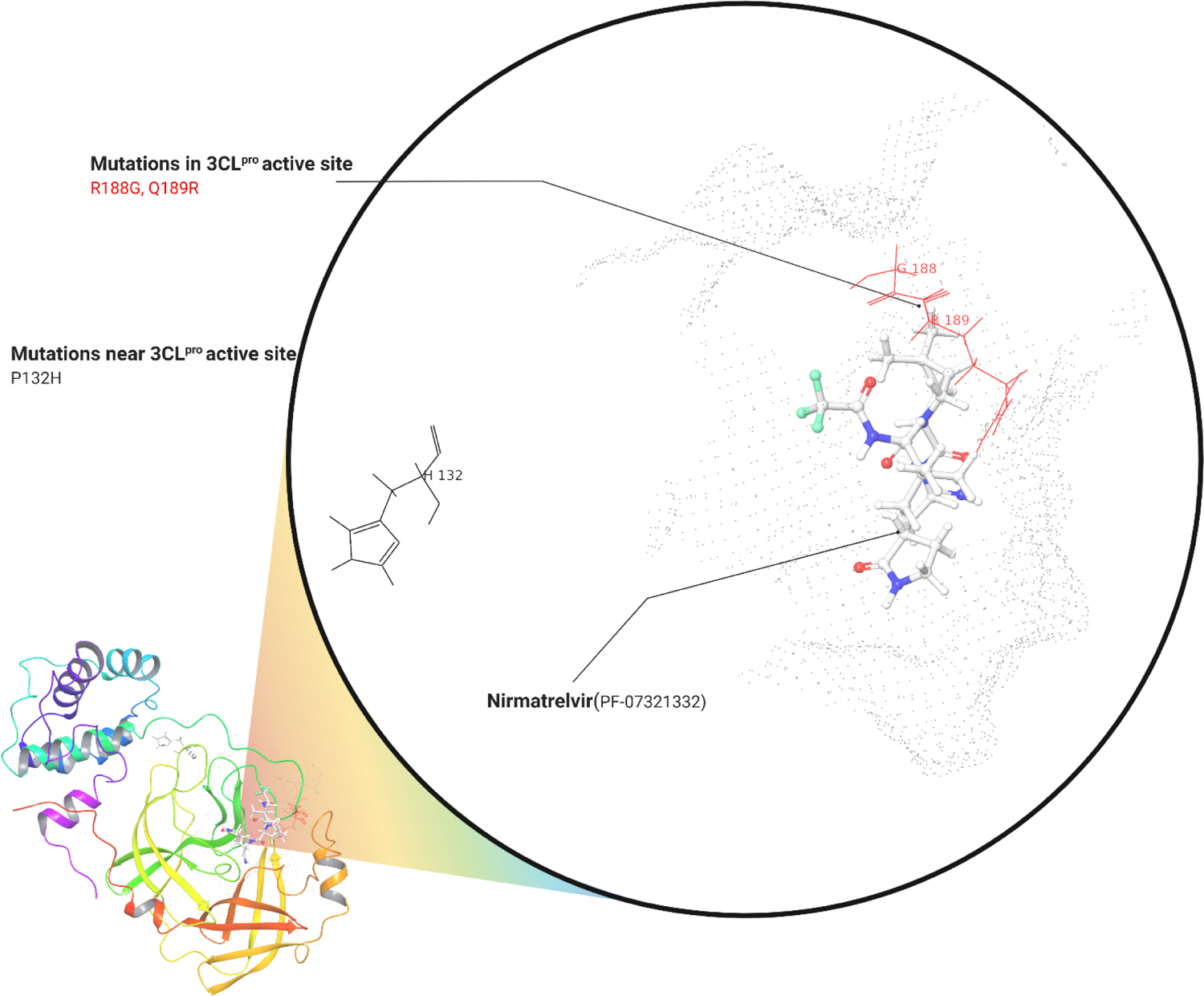
The 3CL-protease Variant3-Italy with grey dots represent the active site region with bind Nirmatrelvir. The red colored sticks represent the mutations in the pocket and the black sticks close to the pocket except the P132H far

**Figure 4D.**
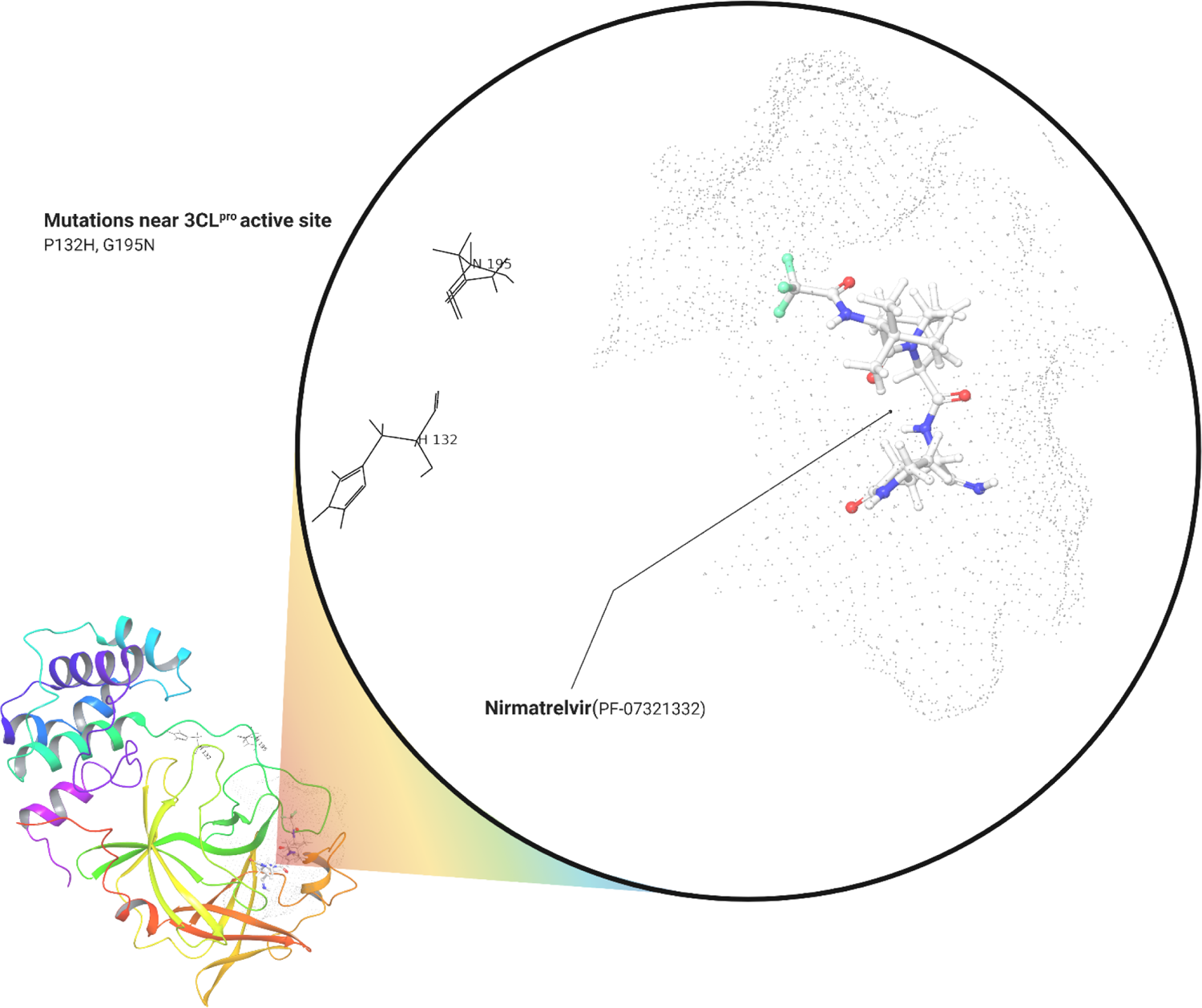
The 3CL-protease Variant4-Italy with grey dots represent the active site region with bind Nirmatrelvir. The black sticks G195N close to the pocket and the P132H far.

### Mutations in the active-site destabilizes the 3CL-protease

Using four different algorithms such as DEUT which also uses two other approaches (mCSM+SDM) and DynaMut2 we predicted the destabilized energies for the Variant1-4 wild type (WT) versus mutant (Mu) in the 3CL-protease residues of the active site and close to pocket which is expressed in change (ΔΔG) energy kcal/mol, the more negative indicates the destability and positive stability. Interestingly, in Variant1-USA the mutation H163W at 163 positions in the active site has stability (ΔΔG) energy of 0.401 kcal/mol (DEUT), 1.36 kcal/mol (DynaMut2) compared to H164N, M165Y and E166Q and residues close to the pocket such as Y161T, M162I, F185L and V186L resulted destability energies in the secondary structure in loop regions of the 3CL-protease in table-2. However, the mutation at P132H 17 Å of distance from the Nirmatrelvir had also destability energy of −1.545 kcal/mol and −1.62 kcal/mol via DEUT and DynaMut2 respectively. In the Variant2-Italy the R188I, Q189G and A191S showed stability of 0.57 kcal/mol, 0.219 kcal/mol and 0.2 kcal/mol to 3CL-protease compared to A194D and G195F destability of −0.1 kcal/mol and −0.63 kcal/mol. The Varian3-4 at Q189R showed stability of 0.35 kcal/mol and R188G and G195N destability of −0.97 kcal/mol and −0.15 kcal/mol respectively. The total (ΔΔG) destibilization energy of the mutant-type (Mut) residues in active site in variant1-USA resulted −3.98 kcal/mol compared to residues outside the pocket −6.8 kcal/mol and in variant2-Italy the total destability energy resulted **-**2.35 kcal/mol for the residues outside the pocket. The variant3-4 resulted total −0.97 kcal/mol and 3.35 kcal/mol for the residues in and outside the pocket respectively.

**Table 2.**
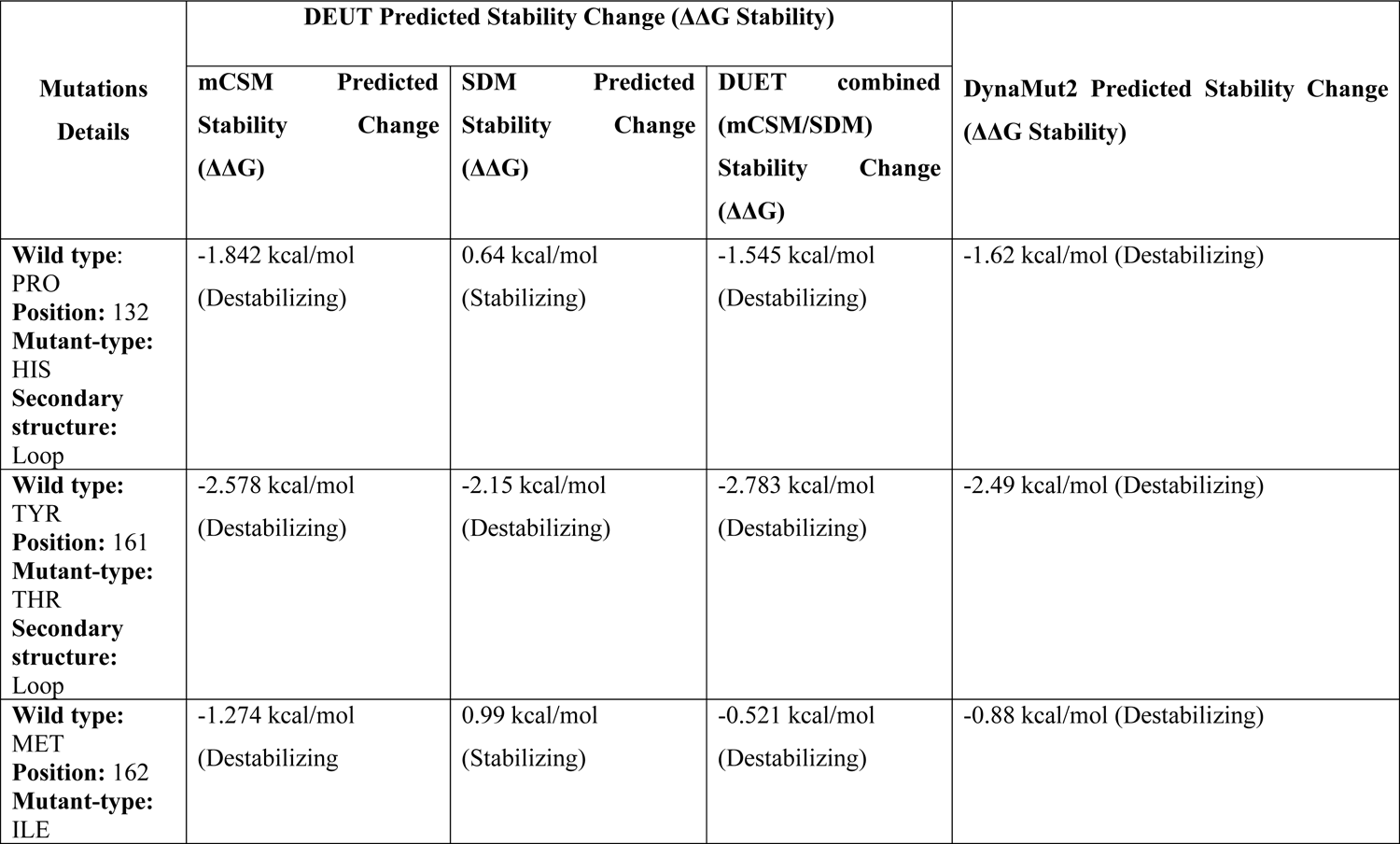
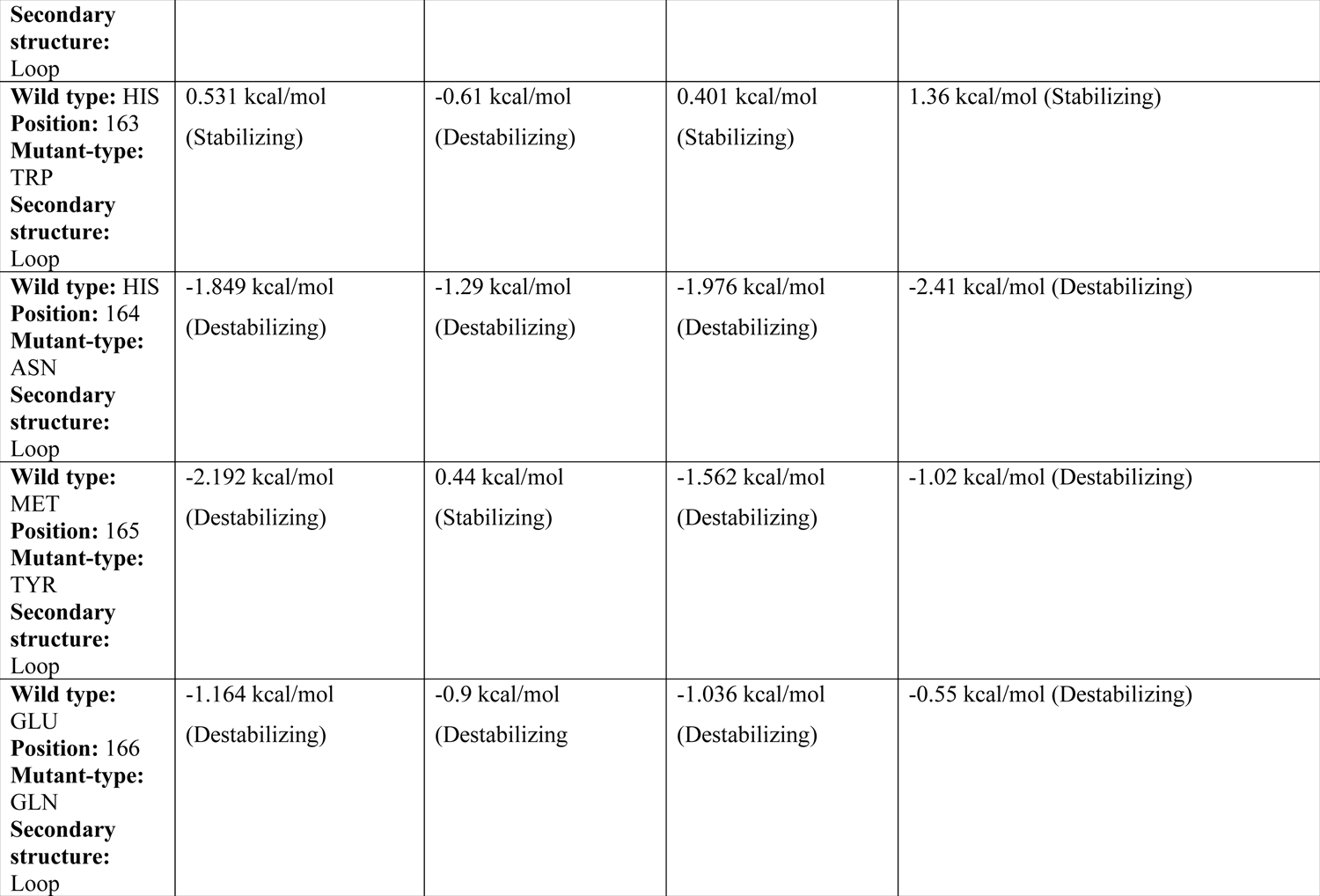
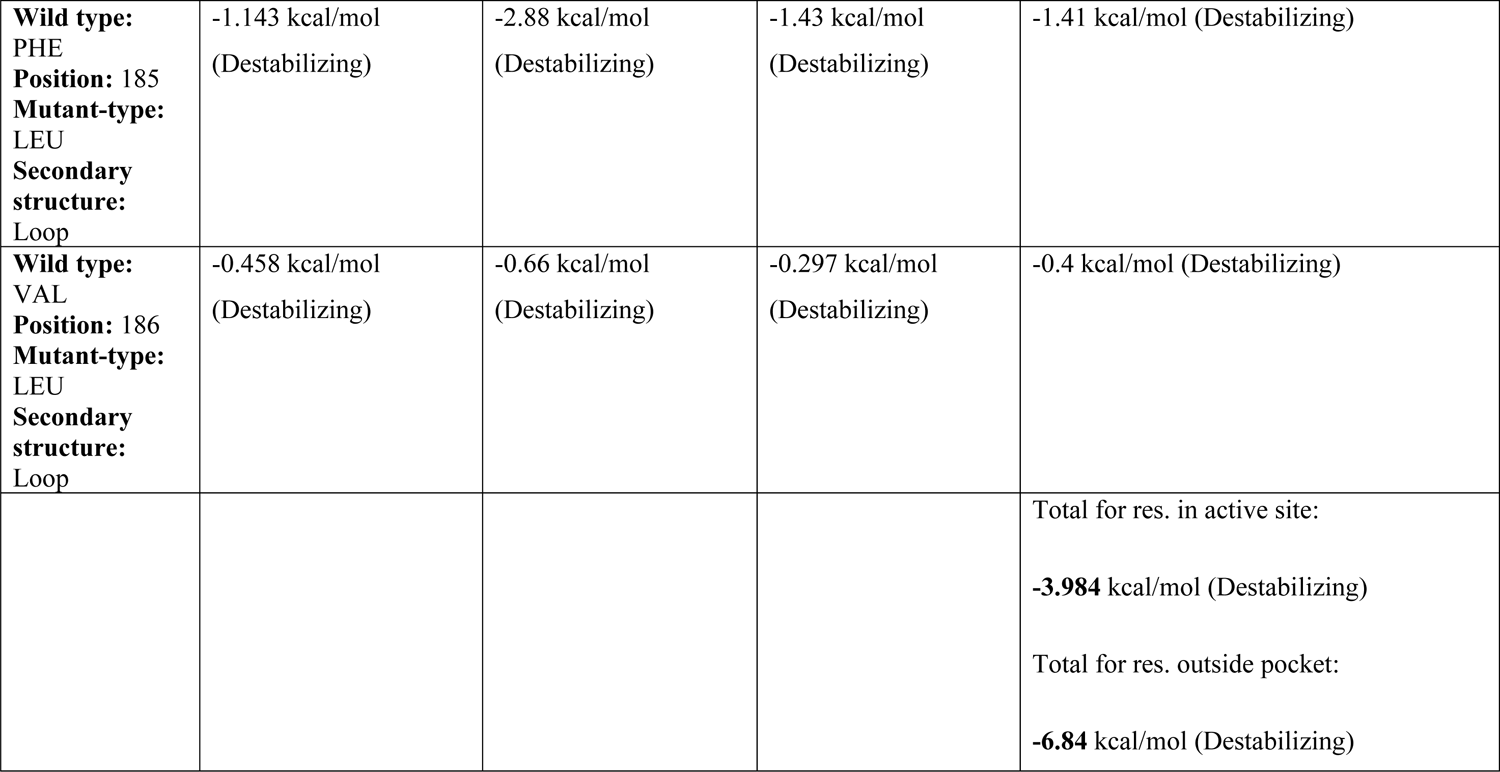
DEUT and DynaMut2 Mutation destabilization analysis of 3CL-protease(**3CL^pro^**) Variant1-USA

**Table 3.** DEUT and DynaMut2 Mutation destabilization analysis of 3CL-protease(**3CL^pro^**) Variant2-Italy

| Mutations Details | DEUT Predicted Stability Change ($\Delta\Delta G$ Stability) | | | DynaMut2 Predicted Stability Change ($\Delta\Delta G$ Stability) |
| --- | --- | --- | --- | --- |
| | mCSM Predicted Stability Change ($\Delta\Delta G$) | SDM Predicted Stability Change ($\Delta\Delta G$) | DUET combined (mCSM+SDM) Predicted Stability Change ($\Delta\Delta G$) | |
| **Wild type:** PRO<br>**Position:** 132<br>**Mutant-type:** HIS<br>**Secondary structure:** Loop | -1.842 kcal/mol (Destabilizing) | 0.64 kcal/mol (Stabilizing) | -1.545 kcal/mol (Destabilizing) | -1.62 kcal/mol (Destabilizing) |
| **Wild type:** ARG<br>**Position:** 188<br>**Mutant-type:** ILE<br>**Secondary structure:** Loop | 0.035 kcal/mol (Stabilizing) | 1.3 kcal/mol (Stabilizing) | 0.498 kcal/mol (Stabilizing) | 0.57 kcal/mol (Stabilizing) |
| **Wild type:** GLN<br>**Position:** 189<br>**Mutant-type:** GLY<br>**Secondary structure:** Loop | -0.33 kcal/mol (Destabilizing) | 1.22 kcal/mol (Stabilizing) | 0.219 kcal/mol (Stabilizing) | -0.18 kcal/mol (Destabilizing) |
| **Wild type:** ALA<br>**Position:** 191<br>**Mutant-type:** SER<br>**Secondary structure:** Loop | -0.698 kcal/mol (Destabilizing) | -1.14 kcal/mol (Destabilizing) | -0.571 kcal/mol (Destabilizing) | 0.2 kcal/mol (Stabilizing) |
| **Wild type:** ALA<br>**Position:** 194<br>**Mutant-type:** ASP | -0.275 kcal/mol (Destabilizing) | -1.79 kcal/mol (Destabilizing) | -0.261 kcal/mol (Destabilizing) | -0.1 kcal/mol (Destabilizing) |
| **Secondary structure:** Loop |  |  |  |  |
| **Wild type:** GLY<br>**Position:** 195<br>**Mutant-type:** PHE<br>**Secondary structure:** Loop | -1.039 kcal/mol<br>(Destabilizing) | -0.14 kcal/mol<br>(Destabilizing) | -0.95 kcal/mol<br>(Destabilizing) | -0.63 kcal/mol (Destabilizing) |
|  |  |  |  | Total for res. outside pocket:<br>**-2.35 kcal/mol (Destabilizing)** |

**Table 4.** DEUT and DynaMut2 Mutation destabilization analysis of 3CL-protease(**3CL^pro^**) Variant3-4 Italy

| **Mutations Details** | **DEUT Predicted Stability Change ($\Delta\Delta G$ Stability)** |  |  | **DynaMut2 Predicted Stability Change ($\Delta\Delta G$ Stability)** |
| --- | --- | --- | --- | --- |
|  | **mCSM Predicted Stability Change ($\Delta\Delta G$)** | **SDM Predicted Stability Change ($\Delta\Delta G$)** | **DUET combined (mCSM+SDM) Predicted Stability Change ($\Delta\Delta G$)** |  |
| **Wild type:** PRO (Variant3-4)<br>**Position:** 132<br>**Mutant-type:** HIS<br>**Secondary structure:** Loop | -1.842 kcal/mol<br>(Destabilizing) | 0.64 kcal/mol<br>(Stabilizing) | -1.545 kcal/mol<br>(Destabilizing) | -1.62 kcal/mol (Destabilizing) |
| **Wild type:** ARG (Variant3)<br>**Position:** 188<br>**Mutant-type:** GLY<br>**Secondary structure:** Loop | -1.186 kcal/mol<br>(Destabilizing) | 0.25 kcal/mol<br>(Stabilizing) | -0.905 kcal/mol<br>(Destabilizing) | -0.97 kcal/mol (Destabilizing) |
| **Wild type:** GLN (Variant3)<br>**Position:** 189<br>**Mutant-type:** ARG<br>**Secondary structure:** Loop | 0.265 kcal/mol<br>(Stabilizing) | 0.47 kcal/mol<br>(Stabilizing) | 0.599 kcal/mol<br>(Stabilizing) | 0.35 kcal/mol (Stabilizing) |
| **Wild type:** GLY (Variant4)<br>**Position:** 195<br>**Mutant-type:** ASN<br>**Secondary structure:** Loop | -0.298 kcal/mol<br>(Destabilizing) | -1.04 kcal/mol<br>(Destabilizing) | -0.175 kcal/mol<br>(Destabilizing) | -0.15 kcal/mol (Destabilizing) |
|  |  |  |  | Total for res. in active site:<br>**-0.97** kcal/mol (Destabilizing)<br>Total for res. outside pocket:<br>**-3.39** kcal/mol (Destabilizing) |

### Impact of mutations on Protein-ligand affinity

Earlier the mutations in the 3CL protease were found to be least such as K90R and G15S in all SARS-CoV-2 variants (DOI: 10.1007/s13238-021-00883-2). However, here we report the presence of several critical mutations in the 3CL protease not reported before making the Omicron VOC a worrisome infectious agent in upcoming days. Because several of these mutations are in the active site region which also destabilizes the 3CL^pro^ thus it is expected that the small molecules could not bind effectively, or their efficacy may reduce to some degree against the 3CL^pro^. Therefore, using the computational technique such as mCSM-lig (https://www.nature.com/articles/srep29575) here we predicted the effect of these critical mutations on protein-ligand binding affinity as well. The Nirmatrelvir binding affinity (Ki = 3.11 nM) was used to compare wild versus mutant type impact of mutations on ligand binding affinity. In the Variant1-USA 3CL^-pro^ revealed that the mutant type residues in and outside the pocket showed the destabilizing affinity change on the ligand binding when compared to wild type and the Y161T stabilizing. The highest affinity change recorded for the M165Y −0.26 log of 3.25 Å distance and lowest the M162I −0.37 log of 9.11 Å distance inside the pocket and outside the pocket to Nirmatrelvir in table 5. And in the Varinat1-Italy Q189G destabilizing −0.90 log 3.38 Å of distance to Nirmatrelvir compared to R188I, A191S, A194D and G195F stabilizing in table 6 and for in the Variant3-4 Q189R −1.108 log destabilizing 3.38 Å of distance to Nirmatrelvir compared to R188G and G195N stabilizing. In all Variant1-4 cases the P132H showed −1.16 log (affinity fold change) of 15.77 Å distance to Nirmatrelvir in table 7.

**Table 5.** mCSM-lig binding affinity analysis of Nirmatrelvir-3CL-protease(**3CL^pro^**) Variant1-USA

| **Mutations Details** | **Ligand ID** | **Distance to ligand** | **Predicted Affinity Change (affinity fold change), Destabilizing in -ve and stabilizing +ve log** |
| --- | --- | --- | --- |
| **Wild type:** PRO<br>**Position:** 132<br>**Mutant-type:** HIS<br>**Secondary structure:** Loop | 7DI | 15.774 Å | -1.168 log |
| **Wild type:** TYR<br>**Position:** 161<br>**Mutant-type:** THR<br>**Secondary structure:** Loop | 7DI | 7.191 Å | 0.366 log |
| **Wild type:** MET<br>**Position:** 162<br>**Mutant-type:** ILE<br>**Secondary structure:** Loop | 7DI | 9.111 Å | -0.374 log |
| **Wild type:** HIS<br>**Position:** 163<br>**Mutant-type:** TRP<br>**Secondary structure:** Loop | 7DI | 2.676 Å | -1.109 log |
| **Wild type:** HIS<br>**Position:** 164<br>**Mutant-type:** ASN<br>**Secondary structure:** Loop | 7DI | 3.017 Å | -1.167 log |
| **Wild type:** MET<br>**Position:** 165<br>**Mutant-type:** TYR<br>**Secondary structure:** Loop | 7DI | 3.251 Å | -2.269 log |
| **Wild type:** GLU<br>**Position:** 166<br>**Mutant-type:** GLN<br>**Secondary structure:** Loop | 7DI | 2.767 Å | -1.817 log |
| **Wild type:** PHE<br>**Position:** 185<br>**Mutant-type:** LEU<br>**Secondary structure:** Loop | 7DI | 6.302 Å | -1.414 log |
| **Wild type:** VAL<br>**Position:** 186<br>**Mutant-type:** LEU<br>**Secondary structure:** Loop | 7DI | 4.588 Å | -1.276 log |

**Table 6.**
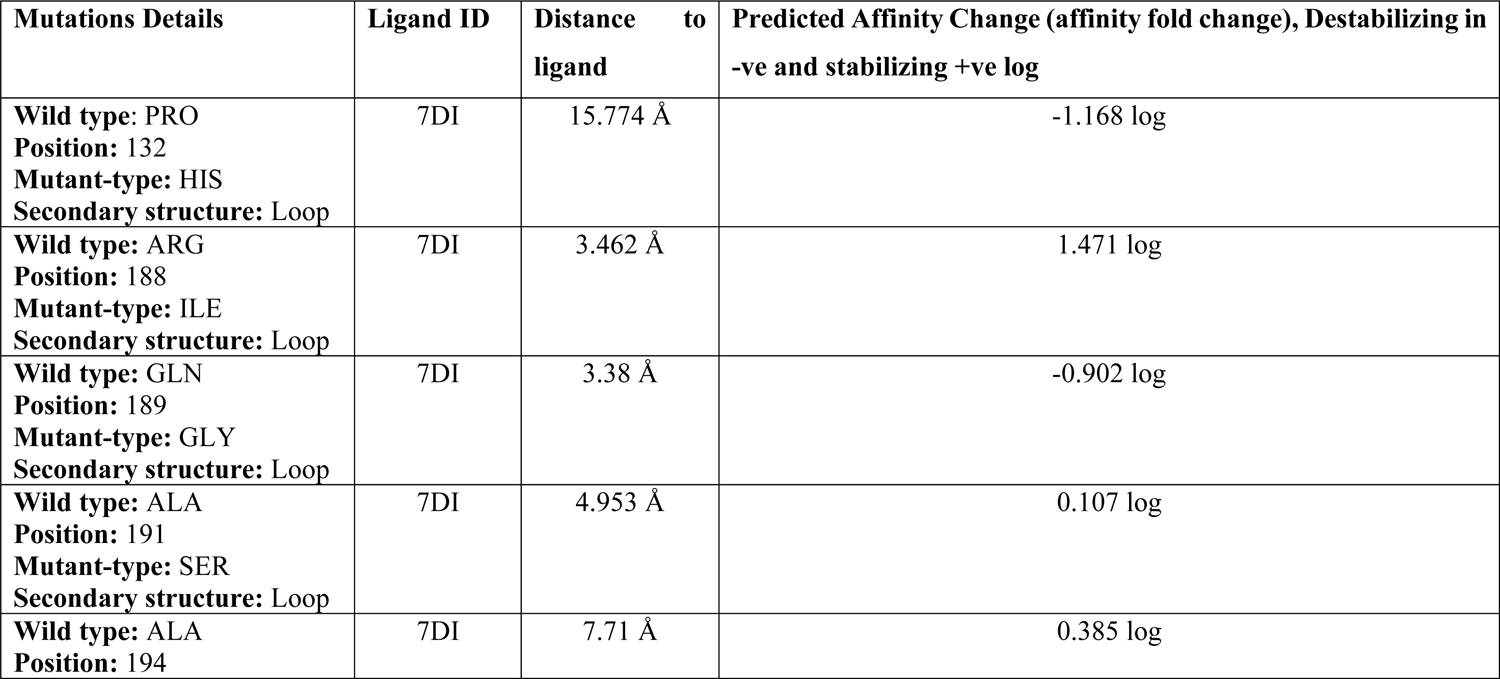
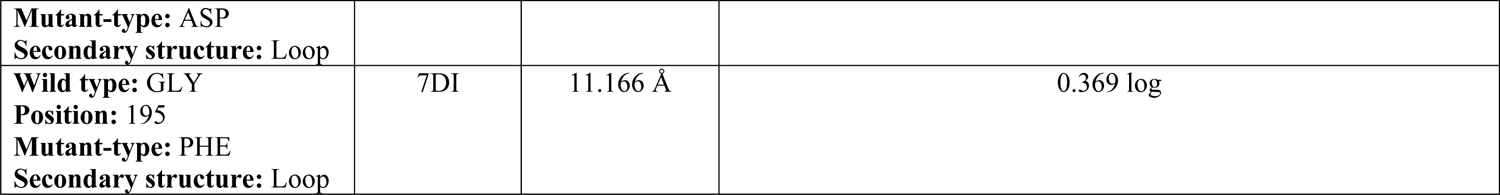
mCSM-lig binding affinity analysis of Nirmatrelvir-3CL-protease (**3CL^pro^**) Variant2-Italy

| **Mutations Details** | **Ligand ID** | **Distance to ligand** | **Predicted Affinity Change (affinity fold change), Destabilizing in -ve and stabilizing +ve log** |
| --- | --- | --- | --- |
| **Wild type:** PRO<br>**Position:** 132<br>**Mutant-type:** HIS<br>**Secondary structure:** Loop | 7DI | 15.774 Å | -1.168 log |
| **Wild type:** ARG<br>**Position:** 188<br>**Mutant-type:** ILE<br>**Secondary structure:** Loop | 7DI | 3.462 Å | 1.471 log |
| **Wild type:** GLN<br>**Position:** 189<br>**Mutant-type:** GLY<br>**Secondary structure:** Loop | 7DI | 3.38 Å | -0.902 log |
| **Wild type:** ALA<br>**Position:** 191<br>**Mutant-type:** SER<br>**Secondary structure:** Loop | 7DI | 4.953 Å | 0.107 log |
| **Wild type:** ALA<br>**Position:** 194 | 7DI | 7.71 Å | 0.385 log |
| **Mutant-type:** ASP<br>**Secondary structure:** Loop |  |  |  |
| **Wild type:** GLY<br>**Position:** 195<br>**Mutant-type:** PHE<br>**Secondary structure:** Loop | 7DI | 11.166 Å | 0.369 log |

**Table 7.** mCSM-lig binding affinity analysis of Nirmatrelvir-3CL-protease (**3CL^pro^**) Variant3-4 Italy

| **Mutations Details** | **Ligand ID** | **Distance to ligand** | **Predicted Affinity Change (affinity fold change), Destabilizing in -ve and stabilizing +ve log** |
| --- | --- | --- | --- |
| **Wild type:** PRO (Variant3-4)<br>**Position:** 132<br>**Mutant-type:** HIS<br>**Secondary structure:** Loop | 7DI | 15.774 Å | -1.168 log |
| **Wild type:** ARG (Variant3)<br>**Position:** 188<br>**Mutant-type:** GLY<br>**Secondary structure:** Loop | 7DI | 3.462 Å | 0.561 log |
| **Wild type:** GLN (Variant3)<br>**Position:** 189<br>**Mutant-type:** ARG<br>**Secondary structure:** Loop | 7DI | 3.38 Å | -1.108 log |
| **Wild type:** GLY (Variant4)<br>**Position:** 195<br>**Mutant-type:** ASN<br>**Secondary structure:** Loop | 7DI | 11.166 Å | 0.419 log |

### Molecular dynamic simulations revealed the CYS145 remained bind to the Nirmatrelvir

Concerned on these several critical mutations inside and close to the binding site of the 3CL^pro^ which may affect the binding pattern of the Nirmatrelvir we also simulated all the Variants1-4 of the 3CL^pro^ and analyzed the binding stability or de-stability pattern and dynamics of Nirmatrelvir whether it remain bind or affected especially of the critical mutations in the binding site in a 100ns simulations. The rmsd(C-α protein) from 0ns-60ns revealed minor fluctuations of rmsd lower than 3.5 Å and onwards to 100ns some major fluctuations of rmsd reached to 3.8 Å and the ligand rmsd(Lig fit on protein) reached to 3.4 Å throughout the 100ns simulations in Variant1-USA complex in figure 5A. The superimposing at 100ns resulted in RMSD of 0.73 Å and no change in the binding pattern of the Nirmatrelvir wild type versus mutant type in figure 5B-C. Moreover, comparing the 2D interactions of the 3CL^pro^-Nirmatrelvir wild type versus mutant type complexes revealed that the covalent bond at CYS145 position remain unaffected at 100ns in figure 5D. To better understand the stability of the CYS145 with Nirmatrelvir, the protein-ligand contacts extracted which revealed a stable CYS145-Nirmatrelvir complex in Variant1-USA throughout the 100ns trajectory in figure 5E. The (C-α protein) rmsd for the Variant2-ITALY revealed much lesser fluctuation of less than 2.2 Å except 2.5 Å rmds at 48ns which then remain stable to onwards 100ns. Similar for the Nirmatrelvir (Lig fit on protein) rmds of less than 2.4 Å except a fluctuation of 2.3 Å in figure 6A. The superimposition of wild versus mutant structures at 100ns frame resulted 2.04 Å rmsd and no change in the binding pattern of the Nirmatrelvir in the binding site of 3CL^pro^ Variant1-ITALY in figure 6B-C. The 2D interactions analysis of the 3CL^pro^-Nirmatrelvir resulted no change in the binding pattern at CYS145 position in figure 6D. Additionally the protein-ligand contact analysis resulted the CYS145 covalent bond interaction with Nirmatrelvir in the binding site of 3CL^pro^ a stable small molecule inhibitor throughout 100ns trajectory in figure 6E.

**Figure 5A.**
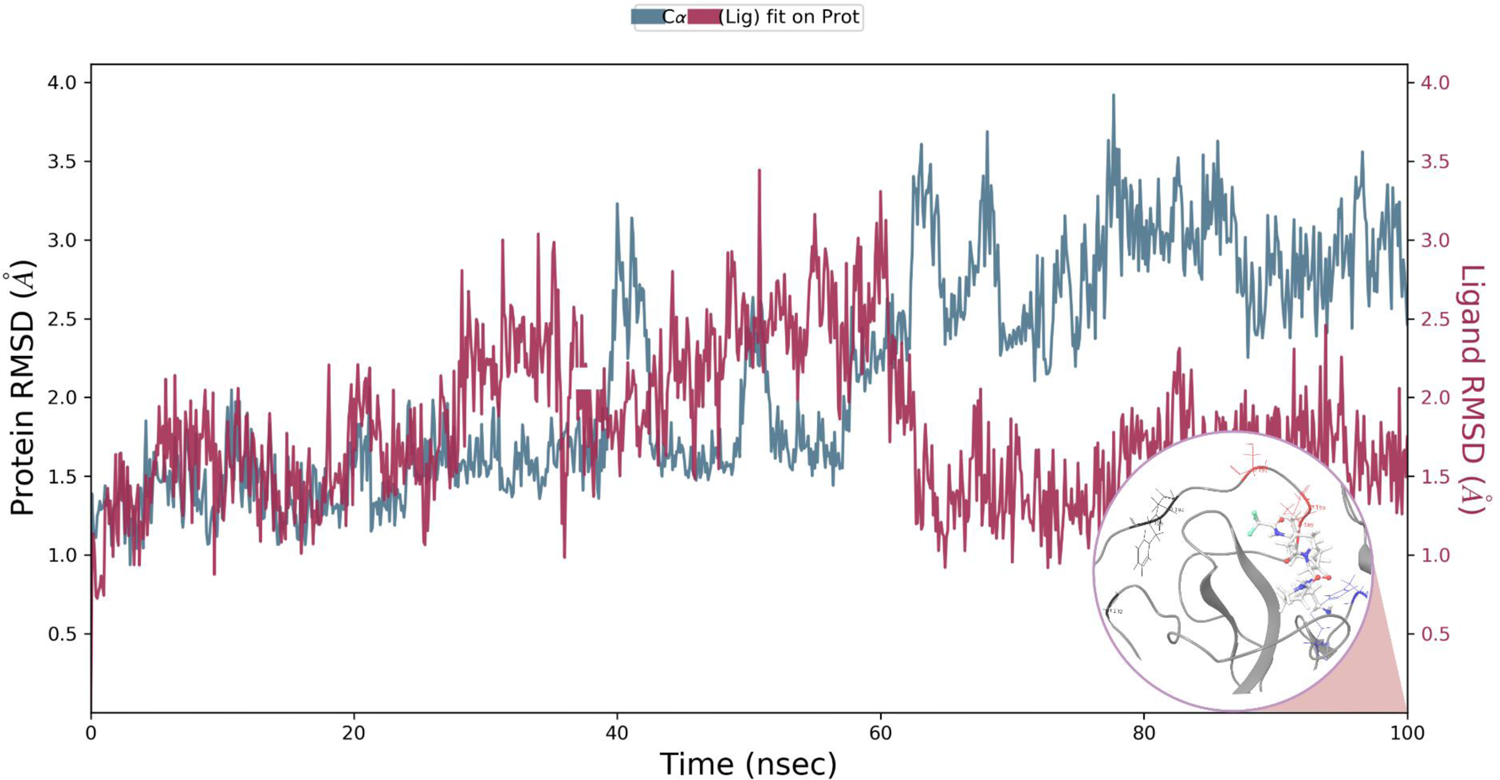
The RMSD analysis of the Variant1-USA 3CL^pro^-Nirmatrelvir complex. The right-hand side scale represents the RMSD of protein(C-α) in teal color and the left ligand in maroon measured in Å. The 3D-complex in grey at 100ns time in the end.

**Figure 5B.**
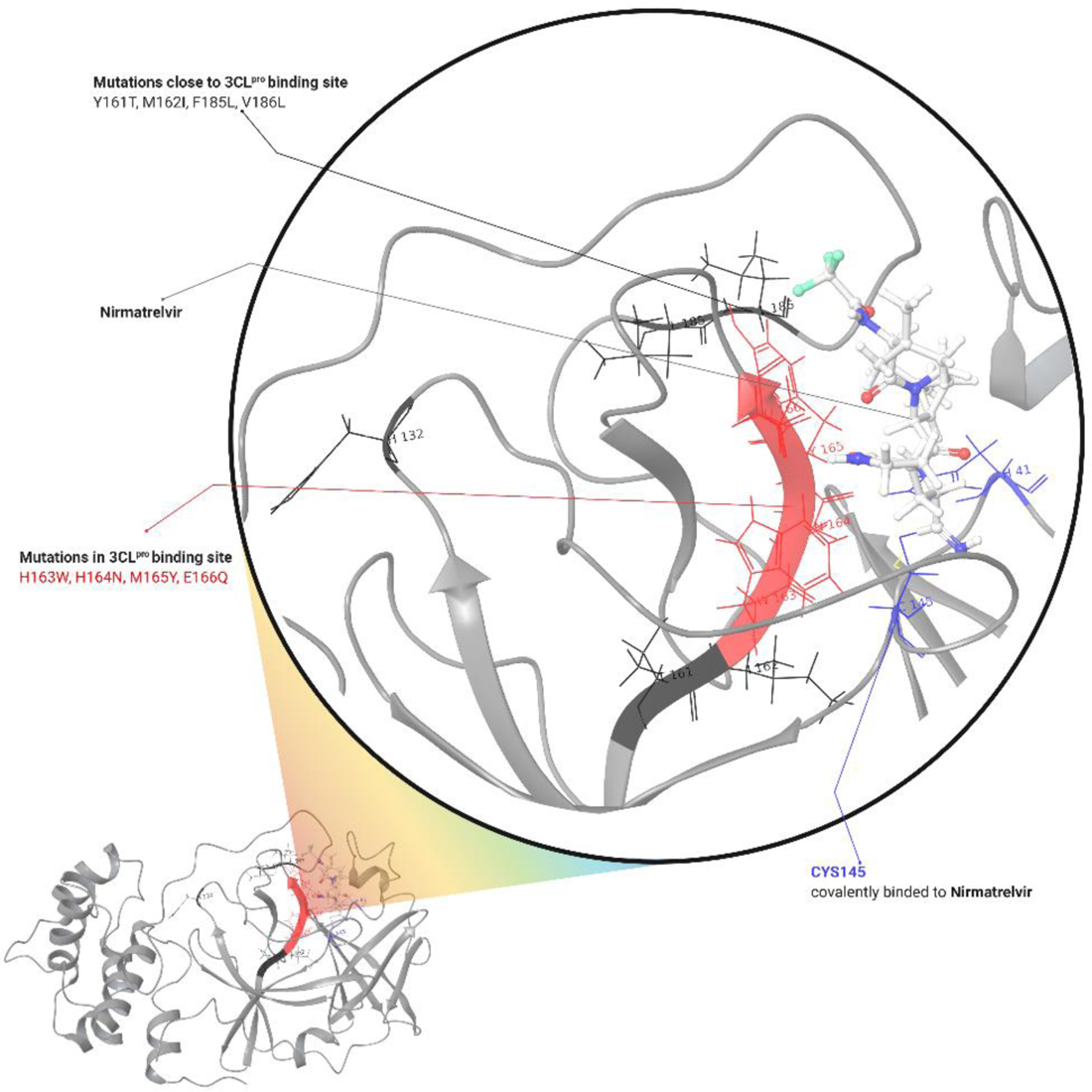
The 3D-structure of the binding site Variant1-USA (3CL^pro^-Nirmatrelvir) at 100ns. The CYS145 in blue, the mutations in the binding site in red color and close to pocket in black and Nirmatrelvir in white ligand representation.

**Figure 5C.**
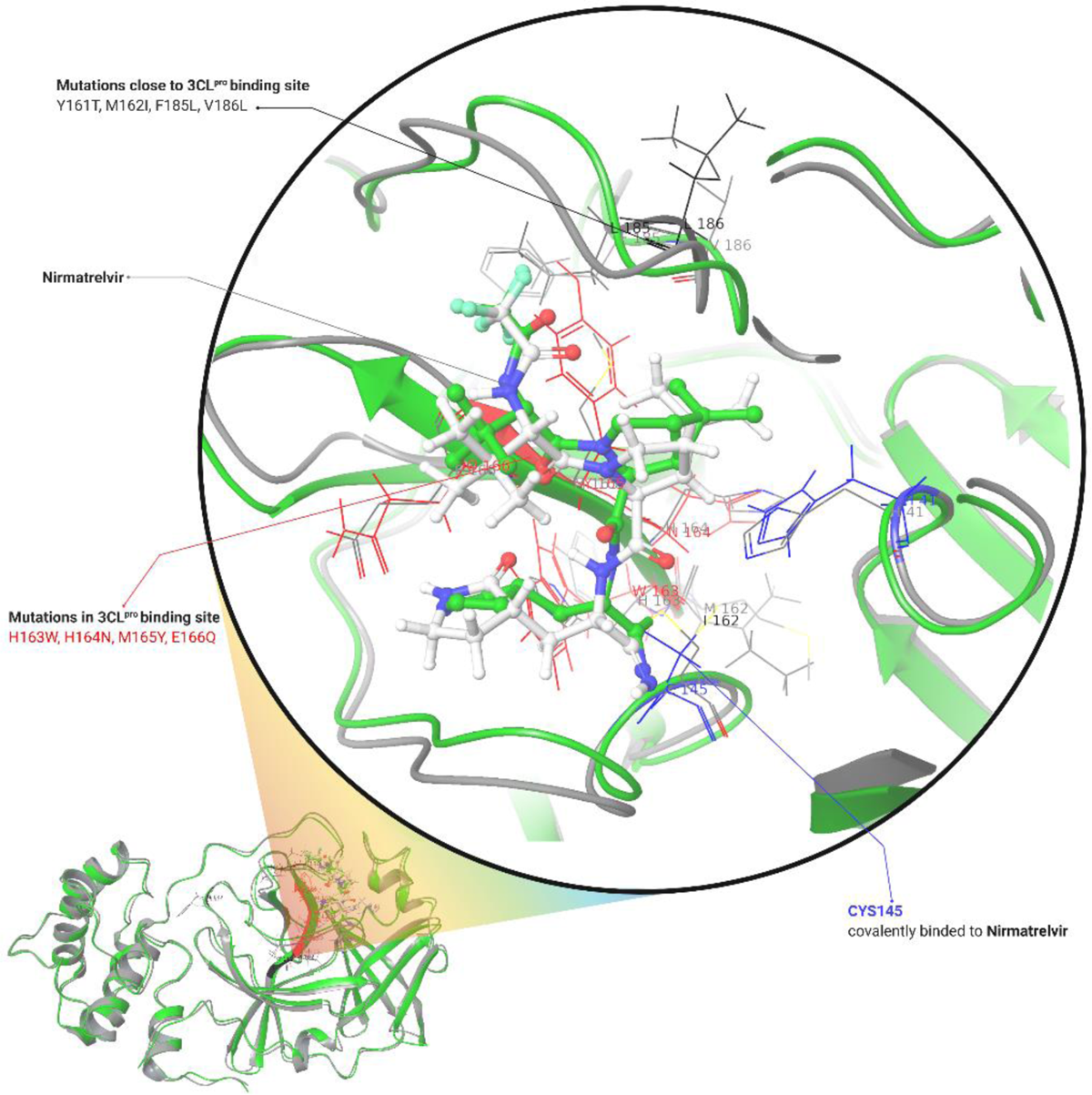
The superpositions of the complex wild (PDB ID:7vh8) versus Variant1-USA in green and grey color respectively. The mutations in the binding site in red and close to pocket in black color. The CYS145 in blue (wild type) and grey (Variant1-USA) remain bonded at 100ns of time.

**Figure 5D.**
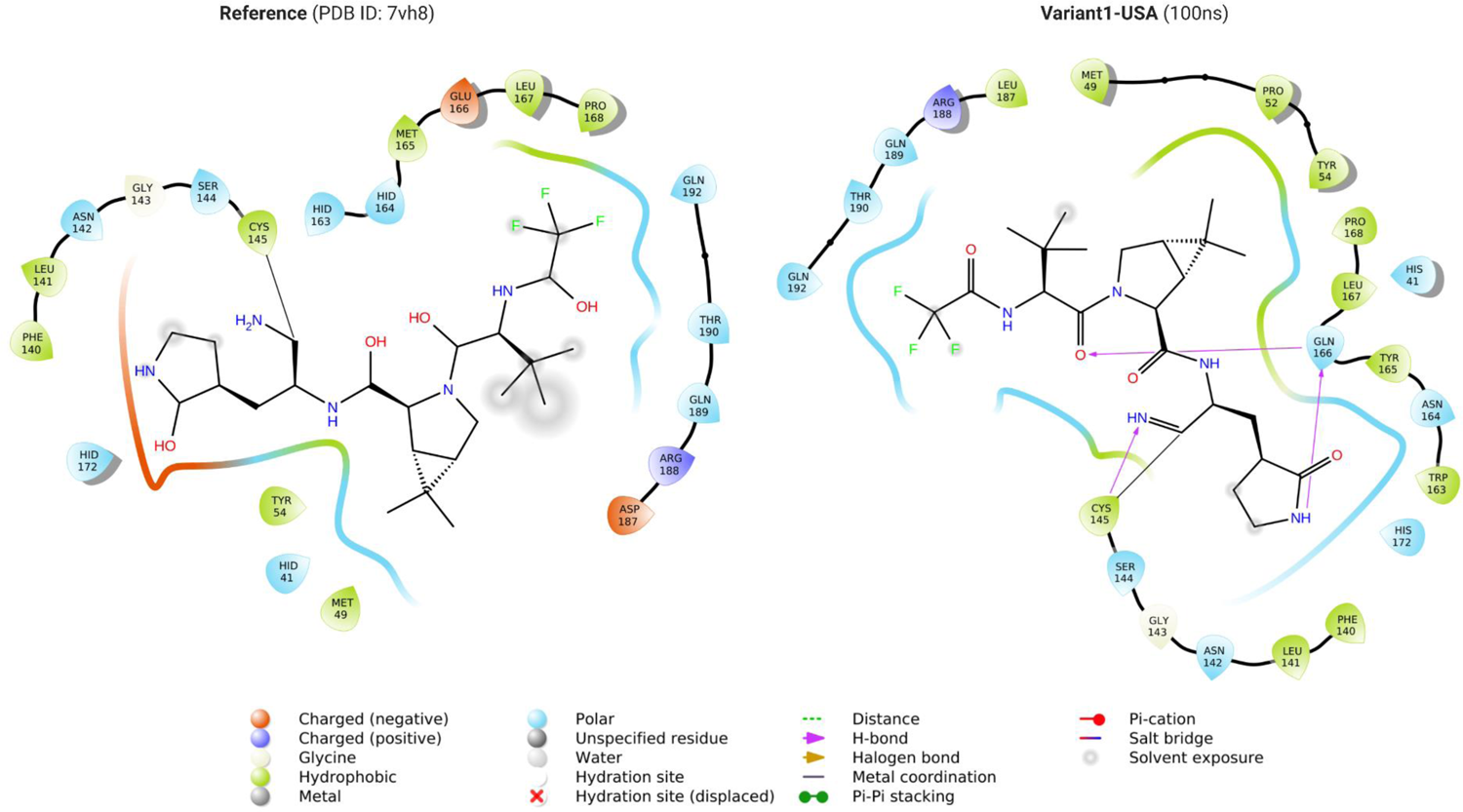
The 2D interactions of the reference structure (PDB ID: 7vh8) versus Variant1-USA at 100ns trajectory. The CYS145 covalent bond interaction represented in black line color remain unaffected Variant1-USA.

**Figure 5E.**
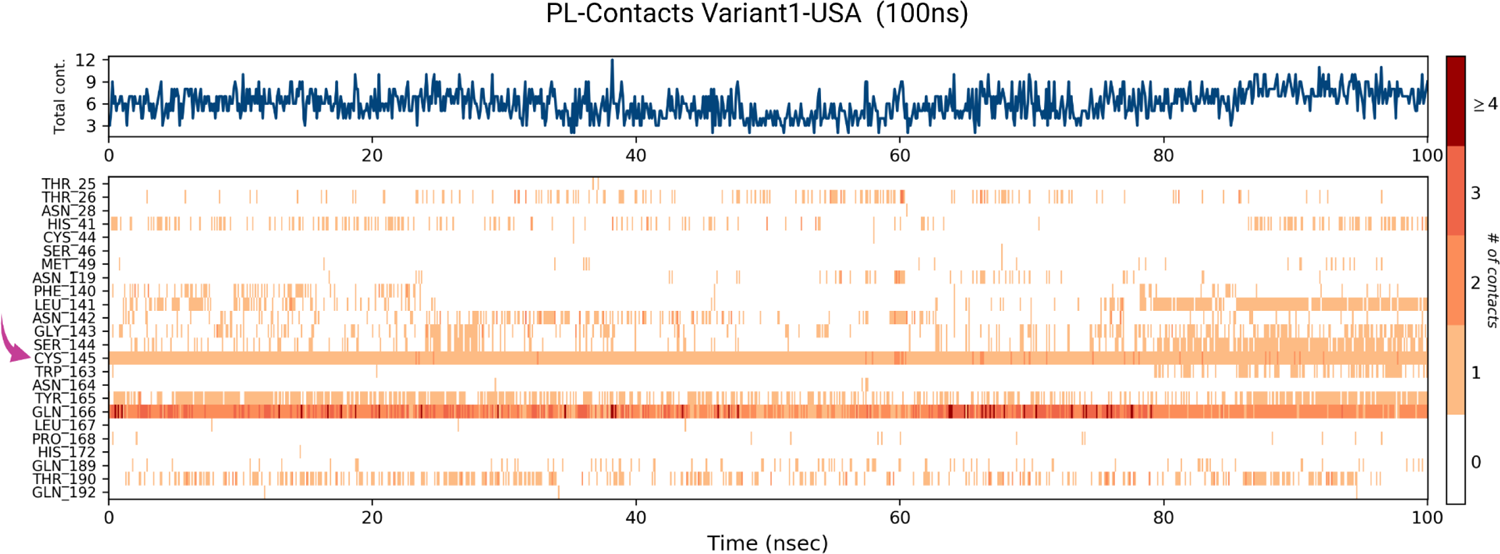
The protein-ligand (PL) contacts analysis of the Variant1-USA (3CL^pro^-Nirmatrelvir) in 100ns trajectory. The CYS145(covalent interaction) highlighted in salmon pink arrow stable (in orange color) in 100ns simulation time.

**Figure 6A.**
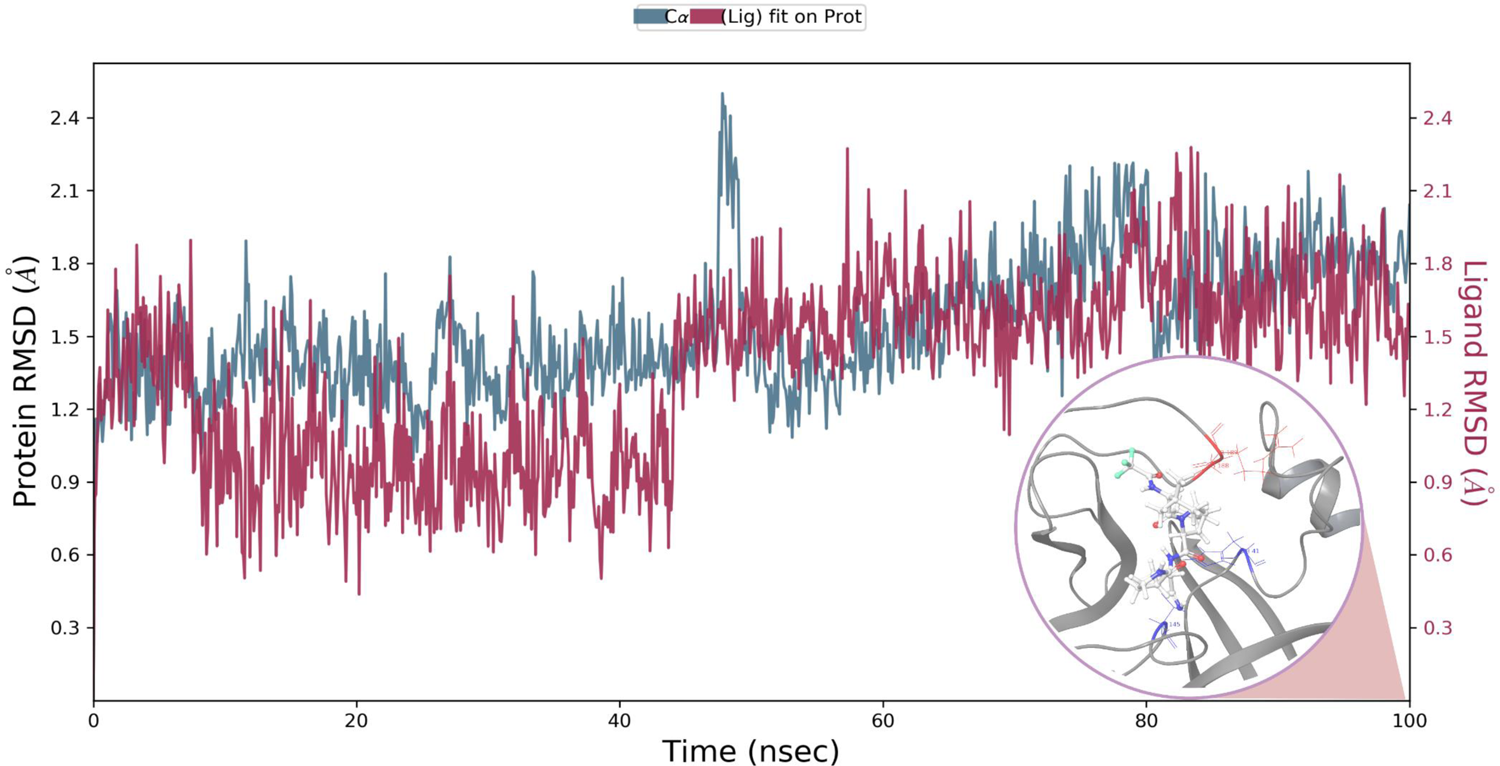
The RMSD analysis of the Variant2-ITALY 3CL^pro^-Nirmatrelvir complex. The right-hand side scale represents the RMSD of protein(C-α) in teal color and the left ligand in maroon measured in Å. The 3D-complex in grey at 100ns time in the end.

**Figure 6B.**
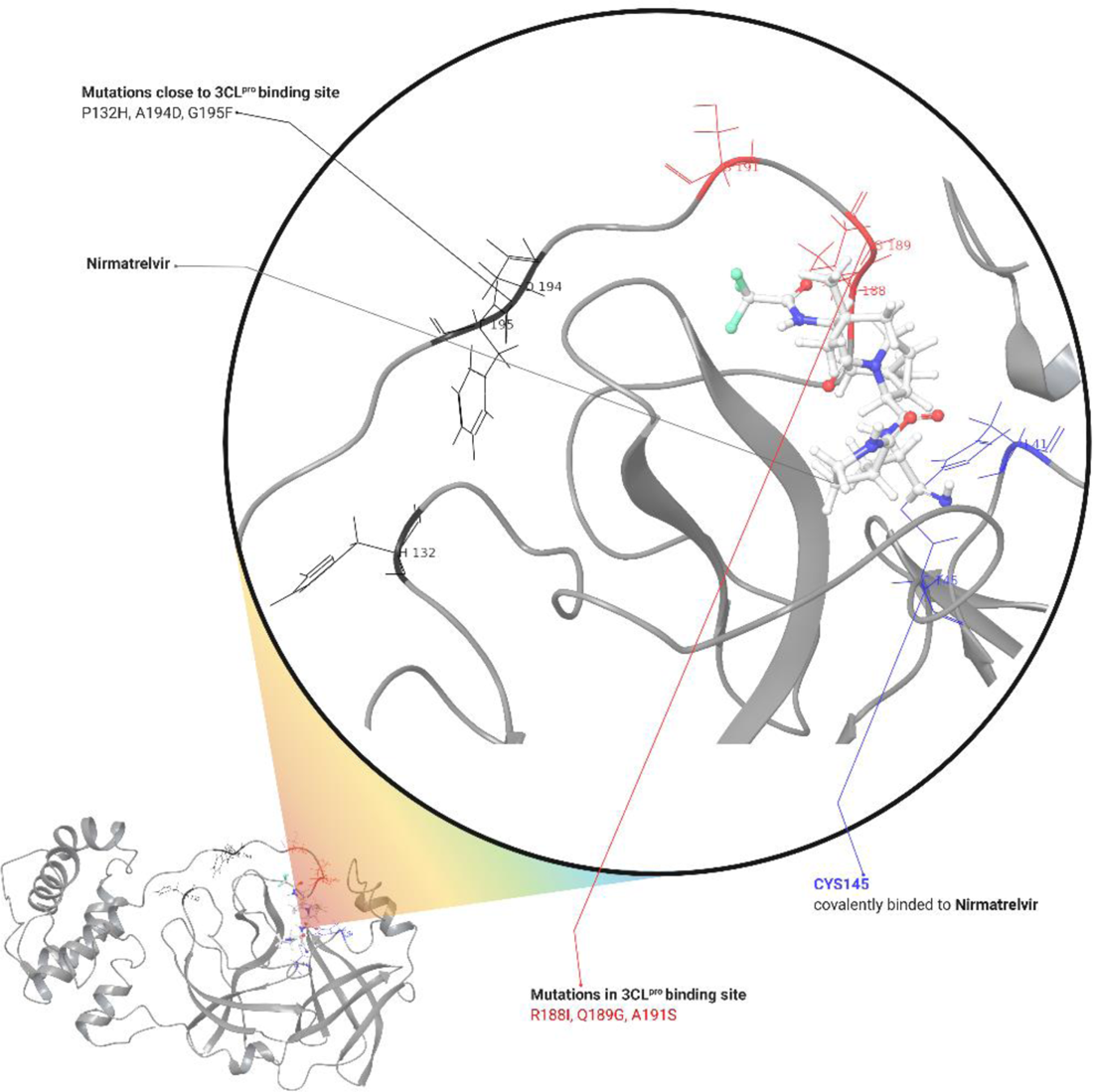
The 3D-structure of the binding site Variant2-ITALY (3CL^pro^-Nirmatrelvir) at 100ns. The CYS145 in blue, the mutations in the binding site in red color and close to pocket in black and Nirmatrelvir in white ligand representation.

**Figure 6C.**
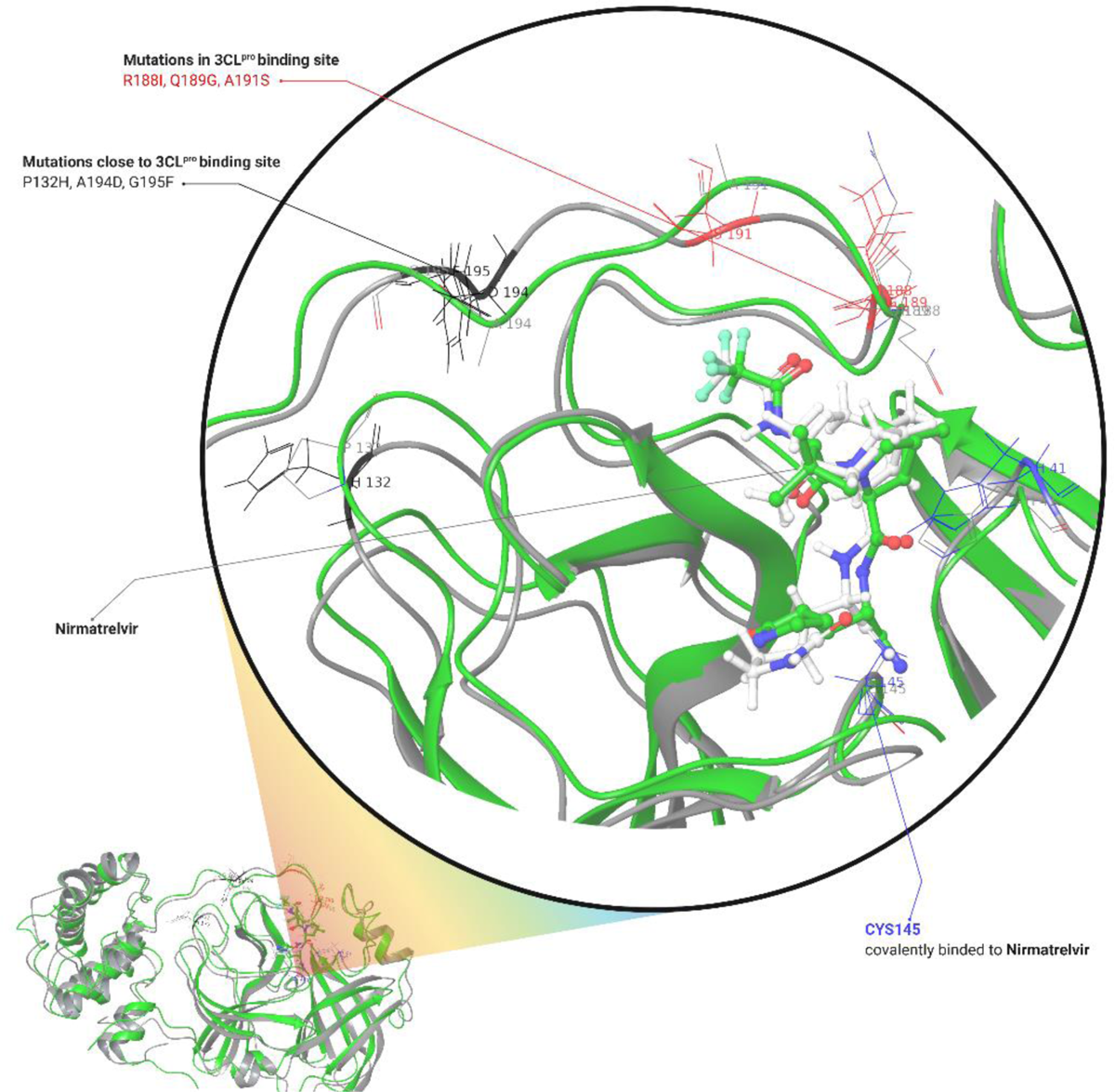
The superposed complex wild (PDB ID:7vh8) versus Variant2-ITALY in green and grey color respectively. The mutations in the binding site in red and close to pocket in black color. The CYS145 in blue (wild type) and grey (Variant2-ITALY) remain bonded at 100ns of time.

**Figure 6D.**
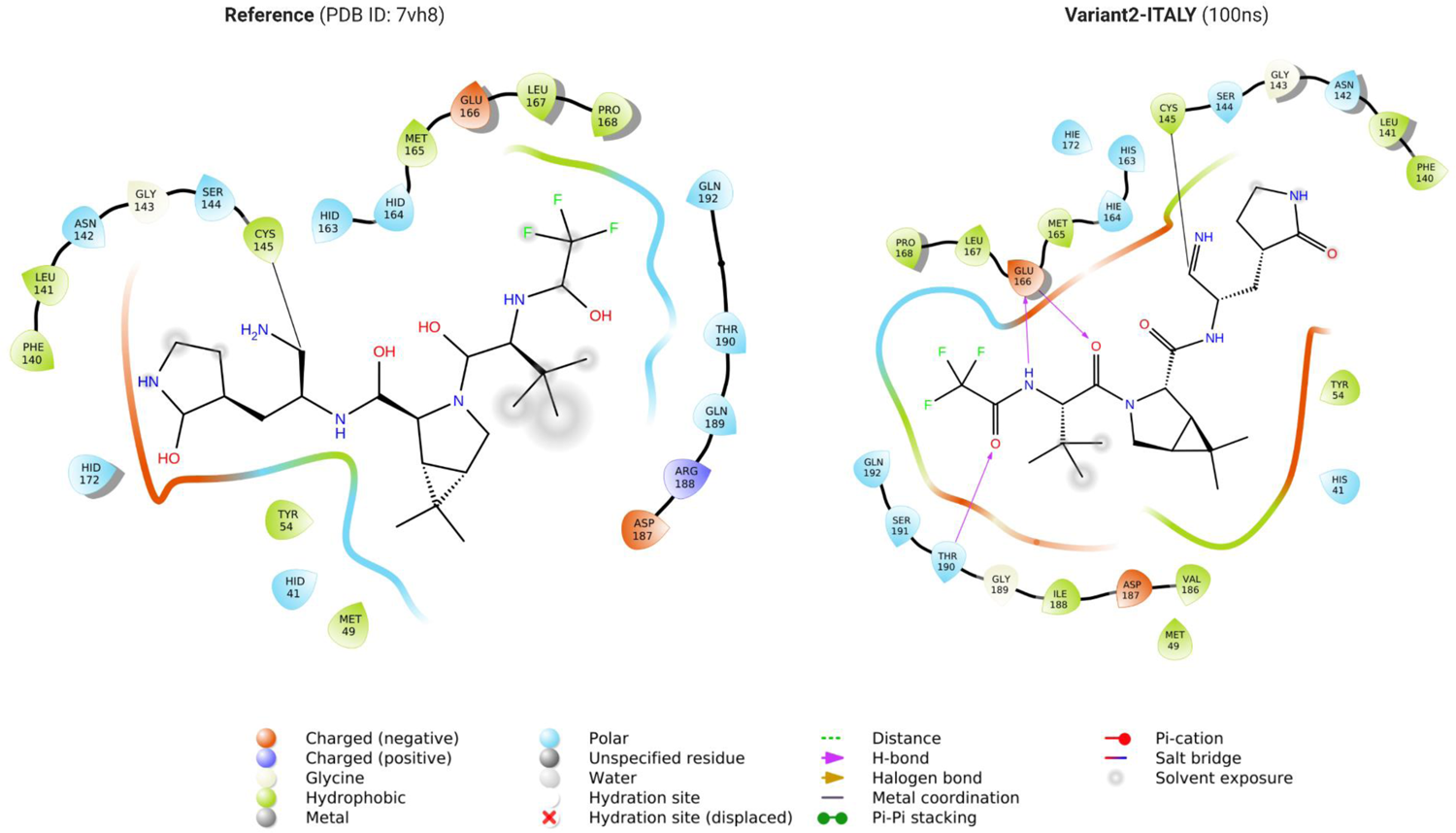
The 2D interactions of the reference structure (PDB ID: 7vh8) versus Variant2-ITALY at 100ns trajectory. The CYS145 covalent bond interaction represented in black line color remained unaffected Variant2-ITALY.

**Figure 6E.**
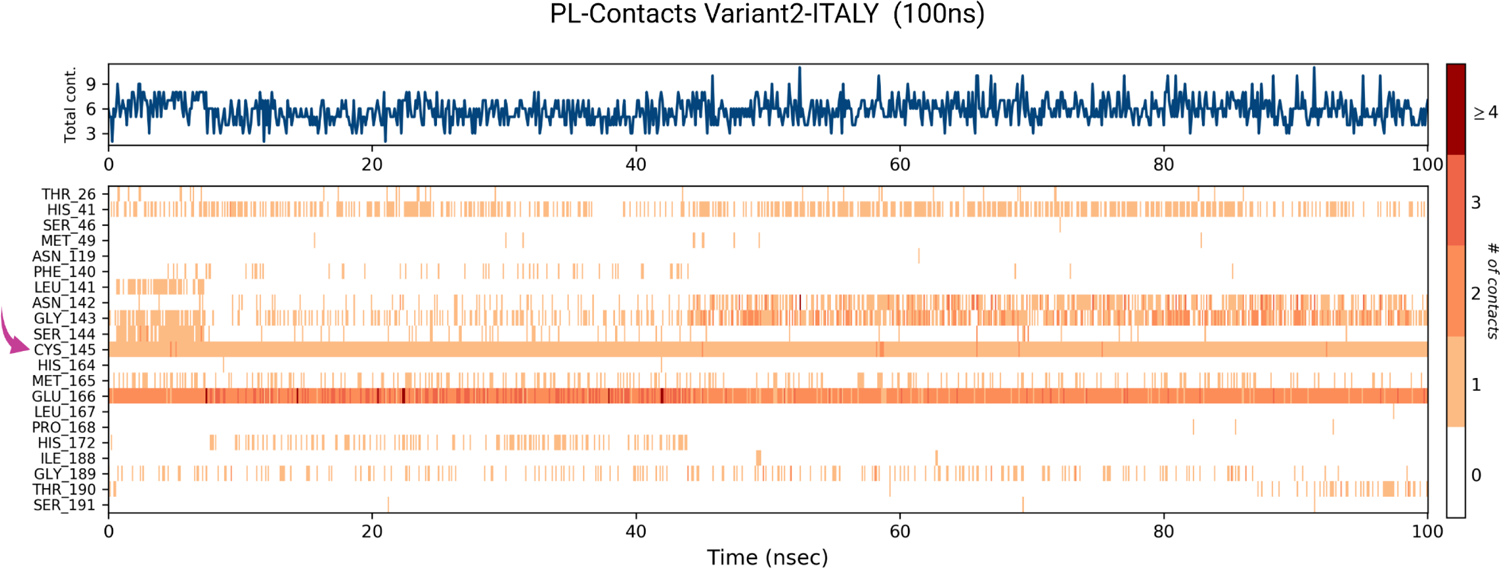
The protein-ligand (PL) contacts analysis of the Variant2-ITALY (3CL^pro^-Nirmatrelvir) in 100ns trajectory. The CYS145(covalent interaction) highlighted in salmon pink arrow stable (in orange color) in 100ns simulation time.

**Figure 7A-B.**
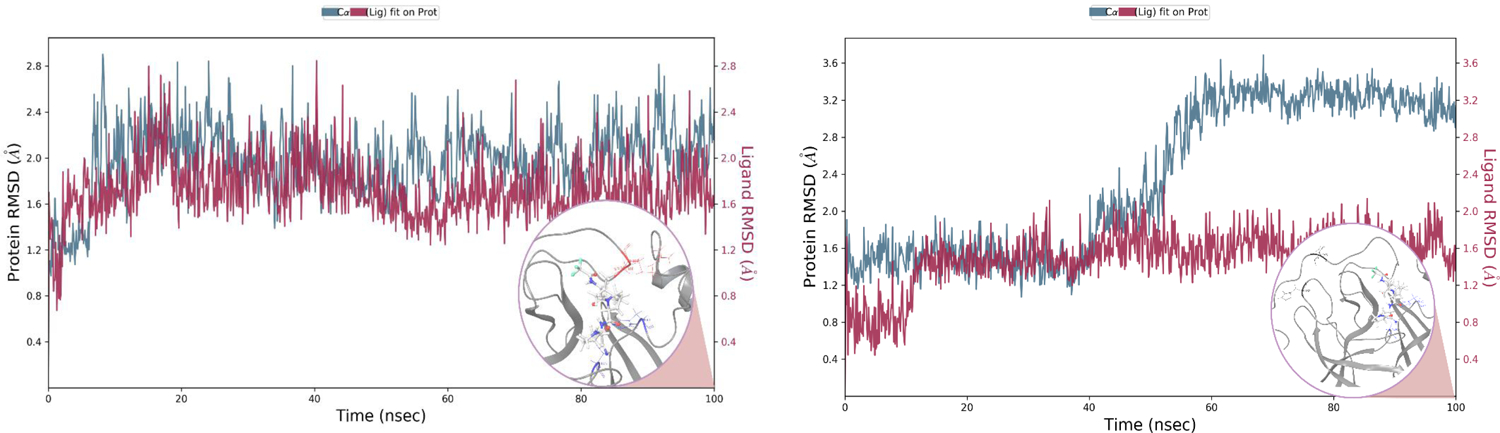
The RMSD analysis of the Variant3-4 ITALY 3CL^pro^-Nirmatrelvir complex. The right-hand side scale represents the RMSD of protein(C-α) in teal color and the left ligand in maroon measured in Å. The 3D-complex in grey at 100ns time in the end.

**Figure 7C-F.**
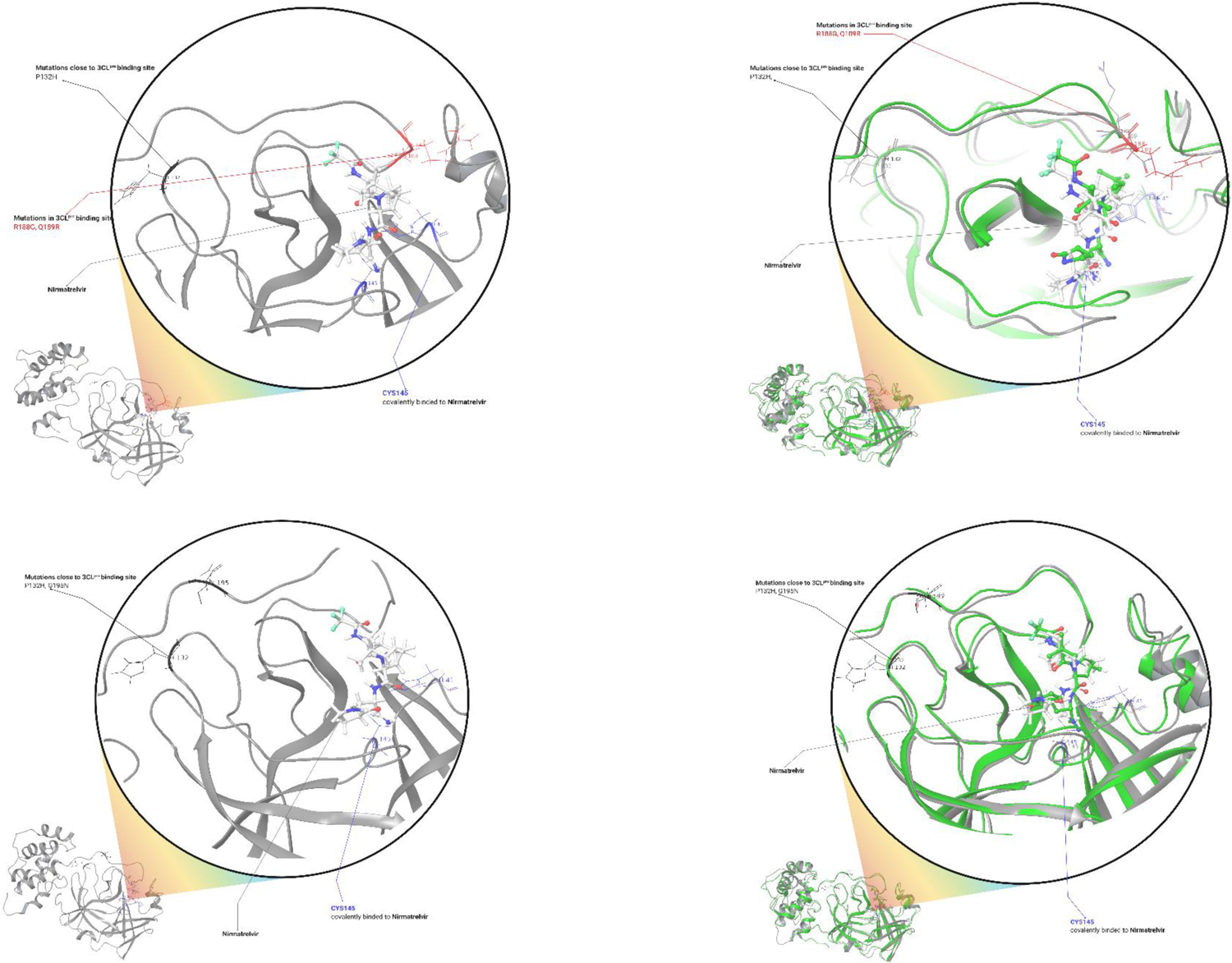
The 2D interactions of the reference structure (PDB ID: 7vh8) versus Variant3-4 ITALY at 100ns trajectory. The CYS145 covalent bond interaction represented in black line color remained unaffected in both cases.

**Figure 7G-H.**
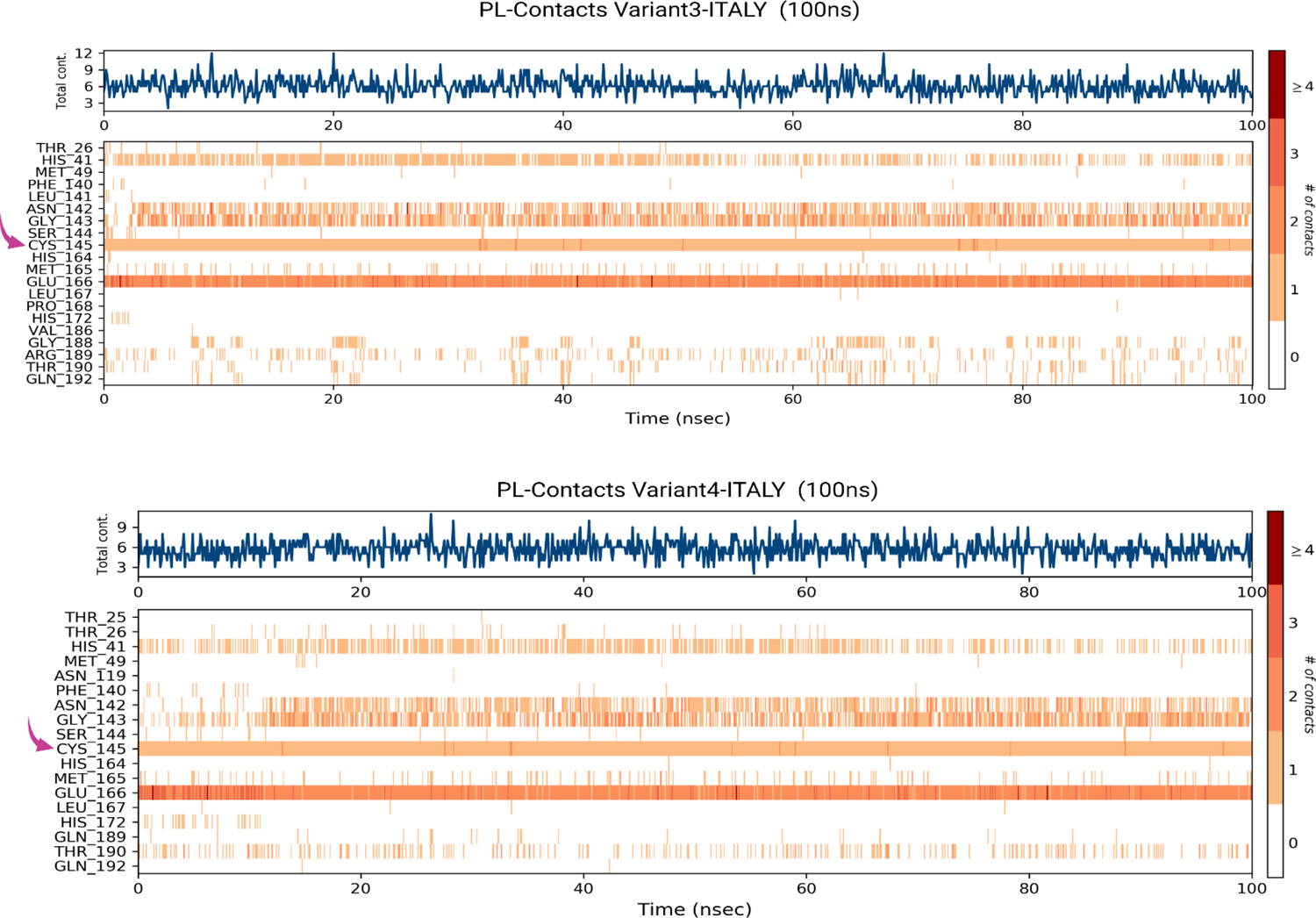
The protein-ligand (PL) contacts analysis of the Variant3-4 ITALY (3CL^pro^-Nirmatrelvir) in 100ns trajectory. The CYS145(covalent interaction) highlighted in salmon pink arrow stable (in orange color) in 100ns simulation time.

## Discussion and Conclusion

In this study, we performed molecular dynamics simulations and binding free energy calculations to investigate the effects of the destabilizing mutations in the binding site of 3CL-protease SARS-CoV-2 Omicron (VOC) on its interaction with its potent inhibitor, Nirmatrelvir an anti-SARS-COV-2 approved oral inhibitor drug molecule. Our results showed that the mutations reduced the binding affinity of Nirmatrelvir by weakening the hydrogen bonds and hydrophobic interactions between the inhibitor and the enzyme. Moreover, the mutations increased the flexibility and conformational entropy of the binding site, leading to a loss of stability and specificity. These findings suggest that the Omicron variant may have a reduced susceptibility to Nirmatrelvir and may be other similar inhibitors that target the 3CL-protease. Therefore, it is important to monitor the emergence and spread of this variant and to develop new strategies to overcome its resistance.

In conclusion, we provided computational insights on the molecular mechanisms underlying the reduced inhibition of 3CL-protease SARS-CoV-2 Omicron by Nirmatrelvir. Our study highlighted the importance of considering the structural and energetic effects of mutations on protein-ligand interactions in drug design and discovery. We hope that our work will contribute to the development of more effective therapeutics against SARS-CoV-2 and its variants.

## Supporting information

Not applicable

## Authors’ contribution

SU designed and leaded the research, wrote the initial draft paper, performed the computational analysis and MD simulations; AA contributed to data analysis and paper preparation; AI guided and contributed in the interpretations of the study; CV and RP contributed to data analysis and paper preparation; GNR contributed to the data interpretation, and wrote parts of the paper.

## References

1 Chathappady House NN, Palissery S, Sebastian H. Corona Viruses: A Review on SARS, MERS and COVID-19. Microbiology Insights 2021; 14: 117863612110024.

2 Roviello V, Roviello GN. Lower COVID-19 mortality in Italian forested areas suggests immunoprotection by Mediterranean plants. Environmental Chemistry Letters 2020.

3 Dai H, Han J, Lichtfouse E. Smarter cures to combat COVID-19 and future pathogens: a review. Environmental Chemistry Letters 2021: 1–13.

4 Bashir MF, Benjiang M, Shahzad L. A brief review of socio-economic and environmental impact of Covid-19. Air Quality, Atmosphere & Health 2020; 13: 1403–9.

5 Caterino M, Gelzo M, Sol S, Fedele R, Annunziata A, Calabrese C, et al. Dysregulation of lipid metabolism and pathological inflammation in patients with COVID-19. Scientific reports 2021; 11: 1–10.

6 Costanzo M, De Giglio MAR, Roviello GN. Anti-Coronavirus Vaccines: Past Investigations on SARS-CoV-1 and MERS-CoV, the Approved Vaccines from BioNTech/Pfizer, Moderna, Oxford/AstraZeneca and others under Development Against SARS-CoV-2 Infection. Current Medicinal Chemistry 2021; 28.

7 He S, Han J, Lichtfouse E. Backward transmission of COVID-19 from humans to animals may propagate reinfections and induce vaccine failure. Environmental Chemistry Letters 2021; 19: 763–8.

8 Zhou D, Dejnirattisai W, Supasa P, Liu C, Mentzer AJ, Ginn HM, et al. Evidence of escape of SARS-CoV-2 variant B.1.351 from natural and vaccine-induced sera. Cell 2021; 184: 2348-61.e6.

9 Costanzo M, De Giglio MA, Roviello GN. SARS-CoV-2: recent reports on antiviral therapies based on lopinavir/ritonavir, darunavir/umifenovir, hydroxychloroquine, remdesivir, favipiravir and other drugs for the treatment of the new coronavirus. Current medicinal chemistry 2020; 27: 4536–41.

10 Borbone N, Piccialli G, Roviello GN, Oliviero G. Nucleoside analogs and nucleoside precursors as drugs in the fight against SARS-CoV-2 and other coronaviruses. Molecules 2021; 26: 986.

11 Kumawat M, Umapathi A, Lichtfouse E, Daima HK. Nanozymes to fight the COVID-19 and future pandemics. Environmental Chemistry Letters 2021.

12 Khan AH, Tirth V, Fawzy M, Mahmoud AED, Khan NA, Ahmed S, et al. COVID-19 transmission, vulnerability, persistence and nanotherapy: a review. Environmental Chemistry Letters 2021; 19: 2773–87.

13 Vicidomini C, Roviello V, Roviello GN. Molecular Basis of the Therapeutical Potential of Clove (Syzygium aromaticum L.) and Clues to Its Anti-COVID-19 Utility. Molecules 2021; 26: 1880.

14 Vicidomini C, Roviello V, Roviello GN. In Silico Investigation on the Interaction of Chiral Phytochemicals from Opuntia ficus-indica with SARS-CoV-2 Mpro. Symmetry 2021; 13: 1041.

15 Wang X, Sun S, Zhang B, Han J. Solar heating to inactivate thermal-sensitive pathogenic microorganisms in vehicles: application to COVID-19. Environmental Chemistry Letters 2020; 19: 1765–72.

16 Yang Y, Islam MS, Wang J, Li Y, Chen X. Traditional Chinese medicine in the treatment of patients infected with 2019-new coronavirus (SARS-CoV-2): a review and perspective. International journal of biological sciences 2020; 16: 1708.

17 Roviello V, Gilhen-Baker M, Vicidomini C, Roviello GN. Forest-bathing and physical activity as weapons against COVID-19: a review. Environmental Chemistry Letters 2021.

18 Roviello V, Roviello GN. Less COVID-19 deaths in southern and insular Italy explained by forest bathing, Mediterranean environment, and antiviral plant volatile organic compounds. Environmental Chemistry Letters 2021.

19 Costanzo M, De Giglio MAR, Roviello GN. SARS CoV-2: Recent Reports on Antiviral Therapies Based on Lopinavir/Ritonavir, Darunavir/Umifenovir, Hydroxychloroquine, Remdesivir, Favipiravir and Other Drugs for the Treatment of the New Coronavirus. Current medicinal chemistry 2020; 27.

20 Bacha U, Barrila J, Velazquez-Campoy A, Leavitt SA, Freire E. Identification of novel inhibitors of the SARS coronavirus main protease 3CLpro. Biochemistry 2004; 43: 4906–12.

21 Modjarrad K. MERS-CoV vaccine candidates in development: The current landscape. Vaccine 2016; 34: 2982–7.

22 Wang L, Shi W, Joyce MG, Modjarrad K, Zhang Y, Leung K, et al. Evaluation of candidate vaccine approaches for MERS-CoV. Nature communications 2015; 6: 1–11.

23 Sui J, Deming M, Rockx B, Liddington RC, Zhu QK, Baric RS, et al. Effects of human anti-spike protein receptor binding domain antibodies on severe acute respiratory syndrome coronavirus neutralization escape and fitness. Journal of virology 2014; 88: 13769–80.

24 Greaney AJ, Loes AN, Crawford KH, Starr TN, Malone KD, Chu HY, et al. Comprehensive mapping of mutations to the SARS-CoV-2 receptor-binding domain that affect recognition by polyclonal human serum antibodies. bioRxiv 2020: 2020.12. 31.425021.

25 Karim SSA, Karim QA. Omicron SARS-CoV-2 variant: a new chapter in the COVID-19 pandemic. The Lancet 2021.

26 Petersen E, Ntoumi F, Hui DS, Abubakar A, Kramer LD, Obiero C, et al. Emergence of new SARS-CoV-2 Variant of Concern Omicron (B.1.1.529) - highlights Africa’s research capabilities, but exposes major knowledge gaps, inequities of vaccine distribution, inadequacies in global COVID-19 response and control efforts. International Journal of Infectious Diseases 2021.

27 Venkatakrishnan A, Anand P, Lenehan PJ, Suratekar R, Raghunathan B, Niesen MJ, et al. Omicron variant of SARS-CoV-2 harbors a unique insertion mutation of putative viral or human genomic origin. 2021.

28 Mahase E. (British Medical Journal Publishing Group, 2021).

29 Shu Y, McCauley J. GISAID: Global initiative on sharing all influenza data – from vision to reality. Eurosurveillance 2017; 22.

30 Zhao Y, Fang C, Zhang Q, Zhang R, Zhao X, Duan Y, et al. Crystal structure of SARS-CoV-2 main protease in complex with protease inhibitor PF-07321332. Protein & Cell 2021.

31 Berman H, Henrick K, Nakamura H. Announcing the worldwide Protein Data Bank. Nature Structural & Molecular Biology 2003; 10: 980-.

32 Madhavi Sastry G, Adzhigirey M, Day T, Annabhimoju R, Sherman W. Protein and ligand preparation: parameters, protocols, and influence on virtual screening enrichments. Journal of Computer-Aided Molecular Design 2013; 27: 221–34.

33 Roos K, Wu C, Damm W, Reboul M, Stevenson JM, Lu C, et al. OPLS3e: Extending Force Field Coverage for Drug-Like Small Molecules. Journal of Chemical Theory and Computation 2019; 15: 1863–74.

34 Harder E, Damm W, Maple J, Wu C, Reboul M, Xiang JY, et al. OPLS3: A Force Field Providing Broad Coverage of Drug-like Small Molecules and Proteins. Journal of Chemical Theory and Computation 2015; 12: 281–96.

35 Bowers KJ, Chow DE, Xu H, Dror RO, Eastwood MP, Gregersen BA, et al. Scalable Algorithms for Molecular Dynamics Simulations on Commodity Clusters. 2006: 43-.

