## Supplementary material for "Computational insights on the destabilizing mutations in the binding site of 3CL-protease SARS-CoV-2 Omicron (VOC)": Not applicable: Supply-Table-S1.pdf

**Table-S1** The 159 Omicron sequences with reported AA (Amino acid) substitutions in the 3CL-protease (3CL$^{pro}$) protein from GISAID.

| 1. S/No. | Accession ID | Location | Gender | Patient age | AA (Amino acid) substitutions in the 3CL-protease (M$^{pro}$) protein sequences |
|---|---|---|---|---|---|
| 2. | EPI_ISL_7496734 | USA / Idaho | Male | 52 | P132H, ins159VT, Y161T, M162I, H163W, H164N, M165Y, E166Q, F185L, V186L, |
| 3. | EPI_ISL_6854347 | Italy / Campania | Female | 8 | P132H, R188I, Q189G, T190stop, A191S, A194D, G195F |
| 4. | EPI_ISL_6854348 | Italy / Campania | Female | 81 | P132H, R188G, Q189R, |
| 5. | EPI_ISL_6854346 | Italy / Campania | Female | 77 | P132H, G195N |
| 6. | EPI_ISL_7192734 | Hong Kong | Male | 37 | P132H |
| 7. | EPI_ISL_7196120 | Nepal / Bagmati | Male | 66 | P132H |
| 8. | EPI_ISL_7196121 | Nepal / Bagmati | Male | 71 | P132H |
| 9. | EPI_ISL_7285845 | South Africa | unknown | unknown | P132H |
| 10. | EPI_ISL_7285846 | South Africa | unknown | unknown | P132H |
| 11. | EPI_ISL_7337497 | SA / Gauteng | Female | 34 | P132H |
| 12. | EPI_ISL_7337502 | SA / Gauteng | Male | 22 | P132H |
| 13. | EPI_ISL_7345322 | UK / England | unknown | unknown | P132H |

| | | | | | |
|---|---|---|---|---|---|
| 14. | EPI_ISL_7381102 | India / New Delhi | Male | 33 | P132H |
| 15. | EPI_ISL_7415767 | USA / Texas | unknown | unknown | P132H |
| 16. | EPI_ISL_7445855 | USA / Iowa | unknown | unknown | P132H |
| 17. | EPI_ISL_7464049 | UK / England | unknown | unknown | P132H |
| 18. | EPI_ISL_7464059 | UK / England | unknown | unknown | P132H |
| 19. | EPI_ISL_7503377 | USA / Arizona | unknown | unknown | P132H |
| 20. | EPI_ISL_7509826 | UK / England | unknown | unknown | P132H |
| 21. | EPI_ISL_7509828 | UK / England | unknown | unknown | P132H |
| 22. | EPI_ISL_7509830 | UK / England | unknown | unknown | P132H |
| 23. | EPI_ISL_7545782 | KwaZulu-Natal | Female | 48 | P132H |
| 24. | EPI_ISL_7552440 | USA / Louisiana | Female | 47 | P132H |
| 25. | EPI_ISL_7552702 | Botswana | Female | 23 | P132H |
| 26. | EPI_ISL_7552703 | Botswana | Female | 44 | P132H |
| 27. | EPI_ISL_7552707 | Botswana | Female | 48 | P132H |
| 28. | EPI_ISL_7563604 | UK / England | unknown | unknown | P132H |
| 29. | EPI_ISL_7566328 | Botswana | Male | 30 | P132H |
| 30. | EPI_ISL_7566330 | Botswana | Female | 30 | P132H |
| 31. | EPI_ISL_7566360 | Botswana | Female | 45 | P132H |
| 32. | EPI_ISL_7566875 | Switzerland | Unknown | Unknown | P132H |
| 33. | EPI_ISL_7583354 | UK | Unknown | Unknown | P132H |

| | | | | | |
|---|---|---|---|---|---|
| 34. | EPI_ISL_7583375 | UK | Unknown | Unknown | P132H |
| 35. | EPI_ISL_7583411 | UK | Unknown | Unknown | P132H |
| 36. | EPI_ISL_7603271 | USA | Unknown | Unknown | P132H |
| 37. | EPI_ISL_7603681 | USA | Unknown | Unknown | P132H |
| 38. | EPI_ISL_7604480 | USA | Unknown | Unknown | P132H |
| 39. | EPI_ISL_7616000 | USA | Female | 43 | P132H |
| 40. | EPI_ISL_7623676 | India | Male | 35 | P132H |
| 41. | EPI_ISL_7625809 | Spain | Male | 32 | P132H |
| 42. | EPI_ISL_7632035 | Switzerland | Unknown | Unknown | P132H |
| 43. | EPI_ISL_7632045 | Switzerland | Unknown | Unknown | P132H |
| 44. | EPI_ISL_7632052 | Switzerland | Unknown | Unknown | P132H |
| 45. | EPI_ISL_7644473 | India | Male | 32 | P132H |
| 46. | EPI_ISL_7649984 | Africa / Namibia | Female | 22 | P132H |
| 47. | EPI_ISL_7649985 | Africa / Namibia | Female | 22 | P132H |
| 48. | EPI_ISL_7649986 | Africa / Namibia | Male | 24 | P132H |
| 49. | EPI_ISL_7649987 | Africa / Namibia | Male | 22 | P132H |
| 50. | EPI_ISL_7649988 | Africa / Namibia | Female | 25 | P132H |
| 51. | EPI_ISL_7649989 | Africa / Namibia | Female | 64 | P132H |
| 52. | EPI_ISL_7649991 | Africa / Namibia | Female | 63 | P132H |
| 53. | EPI_ISL_7649992 | Africa / Namibia | Female | 34 | P132H |

| | | | | | |
|---|---|---|---|---|---|
| 54. | EPI_ISL_7649993 | Africa / Namibia | Female | 54 | P132H |
| 55. | EPI_ISL_7649994 | Africa / Namibia | Female | 22 | P132H |
| 56. | EPI_ISL_7649995 | Africa / Namibia | Male | 48 | P132H |
| 57. | EPI_ISL_7649999 | Africa / Namibia | Female | 32 | P132H |
| 58. | EPI_ISL_7650000 | Africa / Namibia | Male | 37 | P132H |
| 59. | EPI_ISL_7651289 | Lebanon | Male | 49 | P132H |
| 60. | EPI_ISL_7682986 | UK | Unknown | Unknown | P132H |
| 61. | EPI_ISL_7682990 | UK | Unknown | Unknown | P132H |
| 62. | EPI_ISL_7685061 | UK | Unknown | Unknown | P132H |
| 63. | EPI_ISL_7696303 | Belgium | Female | 46 | P132H |
| 64. | EPI_ISL_7701115 | Gauteng | Male | 25 | P132H |
| 65. | EPI_ISL_7701129 | Gauteng | Female | 46 | P132H |
| 66. | EPI_ISL_7703319 | USA | Unknown | Unknown | P132H |
| 67. | EPI_ISL_7703322 | USA | Unknown | Unknown | P132H |
| 68. | EPI_ISL_7725851 | UK | Unknown | Unknown | P132H |
| 69. | EPI_ISL_7725891 | UK | Unknown | Unknown | P132H |
| 70. | EPI_ISL_7725913 | UK | Unknown | Unknown | P132H |
| 71. | EPI_ISL_7725915 | UK | Unknown | Unknown | P132H |
| 72. | EPI_ISL_7725927 | UK | Unknown | Unknown | P132H |
| 73. | EPI_ISL_7733979 | USA | Unknown | Unknown | P132H |

| | | | | | |
|---|---|---|---|---|---|
| 74. | EPI_ISL_7734011 | USA | Unknown | Unknown | P132H |
| 75. | EPI_ISL_7734084 | USA | Unknown | Unknown | P132H |
| 76. | EPI_ISL_7740761 | Gauteng | Female | 23 | P132H |
| 77. | EPI_ISL_7747500 | KwaZulu-Natal | Male | 31 | P132H |
| 78. | EPI_ISL_7747509 | KwaZulu-Natal | Male | 10 | P132H |
| 79. | EPI_ISL_7789703 | Belgium | Male | 62 | P132H |
| 80. | EPI_ISL_7790175 | Belgium | Male | 45 | P132H |
| 81. | EPI_ISL_7799566 | Italy | Male | 25 | P132H |
| 82. | EPI_ISL_7799567 | Italy | Male | 14 | P132H |
| 83. | EPI_ISL_7799568 | Italy | Male | 46 | P132H |
| 84. | EPI_ISL_7799569 | Italy | Male | 14 | P132H |
| 85. | EPI_ISL_7799570 | Italy | Male | 24 | P132H |
| 86. | EPI_ISL_7799571 | Italy | Male | 48 | P132H |
| 87. | EPI_ISL_7799914 | USA | Male | 21 | P132H |
| 88. | EPI_ISL_7799945 | USA | Female | 19 | P132H |
| 89. | EPI_ISL_7830897 | USA | Unknown | Unknown | P132H |
| 90. | EPI_ISL_7834399 | Botswana | Female | 25 | P132H |
| 91. | EPI_ISL_7834407 | Botswana | Male | 11 | P132H |
| 92. | EPI_ISL_7834409 | Botswana | Female | 38 | P132H |
| 93. | EPI_ISL_7834416 | Botswana | Female | 18 | P132H |

| | | | | | |
|---|---|---|---|---|---|
| 94. | EPI_ISL_7834419 | Botswana | Male | 41 | P132H |
| 95. | EPI_ISL_7834422 | Botswana | Female | 34 | P132H |
| 96. | EPI_ISL_7834424 | Botswana | Female | 36 | P132H |
| 97. | EPI_ISL_7834425 | Botswana | Female | 34 | P132H |
| 98. | EPI_ISL_7834426 | Botswana | Female | 21 | P132H |
| 99. | EPI_ISL_7834432 | Botswana | Male | 48 | P132H |
| 100. | EPI_ISL_7834433 | Botswana | Female | 53 | P132H |
| 101. | EPI_ISL_7834434 | Botswana | Male | 39 | P132H |
| 102. | EPI_ISL_7834437 | Botswana | Male | 54 | P132H |
| 103. | EPI_ISL_7834438 | Botswana | Male | 36 | P132H |
| 104. | EPI_ISL_7834439 | Botswana | Male | 33 | P132H |
| 105. | EPI_ISL_7834440 | Botswana | Male | 5 | P132H |
| 106. | EPI_ISL_7834441 | Botswana | Male | 47 | P132H |
| 107. | EPI_ISL_7834442 | Botswana | Female | 38 | P132H |
| 108. | EPI_ISL_7834443 | Botswana | Female | 55 | P132H |
| 109. | EPI_ISL_7834444 | Botswana | Male | 51 | P132H |
| 110. | EPI_ISL_7834446 | Botswana | Male | 34 | P132H |
| 111. | EPI_ISL_7834447 | Botswana | Male | 25 | P132H |
| 112. | EPI_ISL_7834449 | Botswana | Male | 56 | P132H |
| 113. | EPI_ISL_7834451 | Botswana | Female | 16 | P132H |

| | | | | | |
|---|---|---|---|---|---|
| 114. | EPI_ISL_7834452 | Botswana | Male | 19 | P132H |
| 115. | EPI_ISL_7834453 | Botswana | Female | 24 | P132H |
| 116. | EPI_ISL_7834455 | Botswana | Male | 24 | P132H |
| 117. | EPI_ISL_7834456 | Botswana | Female | 36 | P132H |
| 118. | EPI_ISL_7834457 | Botswana | Female | 35 | P132H |
| 119. | EPI_ISL_7834458 | Botswana | Female | 4 | P132H |
| 120. | EPI_ISL_7834459 | Botswana | Male | 32 | P132H |
| 121. | EPI_ISL_7834461 | Botswana | Male | 31 | P132H |
| 122. | EPI_ISL_7834462 | Botswana | Male | 55 | P132H |
| 123. | EPI_ISL_7834463 | Botswana | Male | 40 | P132H |
| 124. | EPI_ISL_7834465 | Botswana | Male | 29 | P132H |
| 125. | EPI_ISL_7834466 | Botswana | Male | 22 | P132H |
| 126. | EPI_ISL_7834467 | Botswana | Male | 8 | P132H |
| 127. | EPI_ISL_7834468 | Botswana | Male | 60 | P132H |
| 128. | EPI_ISL_7834472 | Botswana | Female | 44 | P132H |
| 129. | EPI_ISL_7834473 | Botswana | Female | 6 days | P132H |
| 130. | EPI_ISL_7834475 | Botswana | Female | 15 | P132H |
| 131. | EPI_ISL_7834476 | Botswana | Male | 20 | P132H |
| 132. | EPI_ISL_7834477 | Botswana | Male | 2 | P132H |
| 133. | EPI_ISL_7834480 | Botswana | Male | 39 | P132H |

| | | | | | |
|---|---|---|---|---|---|
| 134. | EPI_ISL_7834481 | Botswana | Male | 4 | P132H |
| 135. | EPI_ISL_7834482 | Botswana | Male | 36 | P132H |
| 136. | EPI_ISL_7834493 | Botswana | Female | 49 | P132H |
| 137. | EPI_ISL_7834502 | Botswana | Male | 27 | P132H |
| 138. | EPI_ISL_7834514 | Botswana | Male | 15 | P132H |
| 139. | EPI_ISL_7834554 | Botswana | Male | 38 | P132H |
| 140. | EPI_ISL_7834555 | Botswana | Female | 35 | P132H |
| 141. | EPI_ISL_7834572 | Botswana | Male | 42 | P132H |
| 142. | EPI_ISL_7834590 | Botswana | Male | 31 | P132H |
| 143. | EPI_ISL_7834591 | Botswana | Female | 17 | P132H |
| 144. | EPI_ISL_7834600 | Botswana | Male | 9 | P132H |
| 145. | EPI_ISL_7843337 | UK | Unknown | Unknown | P132H |
| 146. | EPI_ISL_7843349 | UK | Unknown | Unknown | P132H |
| 147. | EPI_ISL_7843355 | UK | Unknown | Unknown | P132H |
| 148. | EPI_ISL_7862534 | UK | Unknown | Unknown | P132H |
| 149. | EPI_ISL_7864703 | India | Female | 24 | P132H |
| 150. | EPI_ISL_7876997 | India | Female | 20 | P132H |
| 151. | EPI_ISL_7877006 | India | Male | 19 | P132H |
| 152. | EPI_ISL_7877093 | India | Female | 39 | P132H |
| 153. | EPI_ISL_7877115 | India | Male | 6 | P132H |

| | | | | | |
|---|---|---|---|---|---|
| 154. | EPI_ISL_7877191 | India | Male | 40 | P132H |
| 155. | EPI_ISL_7877201 | India | Male | 18 | P132H |
| 156. | EPI_ISL_7877202 | India | Male | 57 | P132H |
| 157. | EPI_ISL_7877203 | India | Female | 19 | P132H |
| 158. | EPI_ISL_7877297 | India | Male | 23 | P132H |
| 159. | EPI_ISL_7887528 | USA | Unknown | Unknown | P132H |
